## Supplementary material for "Analysing and simulating energy-based models in biology using BondGraphTools": LaTeX source for manuscript: bibliography.bib

@article{Soh2010Network,
	author = {Soh, Keng Cher and Hatzimanikatis, Vassily},
	title = {Network thermodynamics in the post-genomic era},
	journal = {Current Opinion in Microbiology},
	volume = {13},
	number = {3},
	pages = {350--357},
	year = {2010},
	month = {Jun},
	issn = {1369-5274},
	publisher = {Elsevier Current Trends},
	doi = {10.1016/j.mib.2010.03.001}
}

@article{Flamholz2012Jan,
    author = {Flamholz, Avi and Noor, Elad and Bar-Even, Arren and Milo, Ron},
    title = {eQuilibrator--the biochemical thermodynamics calculator},
    journal = {Nucleic Acids Research},
    volume = {40},
    number = {D1},
    pages = {D770--D775},
    year = {2012},
    month = {Jan},
    issn = {0305-1048},
    publisher = {Oxford Academic},
    doi = {10.1093/nar/gkr874}
}

@article{Oster1973Network,
	author = {Oster, George F. and Perelson, Alan S. and Katchalsky, Aharon},
	title = {Network thermodynamics: dynamic modelling of biophysical systems},
	journal = {Quarterly Reviews of Biophysics},
	volume = {6},
	number = {1},
	pages = {1--134},
	year = {1973},
	month = {Feb},
	issn = {1469-8994},
	publisher = {Cambridge University Press},
	doi = {10.1017/S0033583500000081}
}

@article{Ederer2007Thermodynamically,
	author = {Ederer, Michael and Gilles, Ernst Dieter},
	title = {Thermodynamically Feasible Kinetic Models of Reaction Networks},
	journal = {Biophysical Journal},
	volume = {92},
	number = {6},
	pages = {1846--1857},
	year = {2007},
	month = {Mar},
	issn = {0006-3495},
	publisher = {Elsevier},
	doi = {10.1529/biophysj.106.094094}
}

@article{Liebermeister2006Bringing,
	author = {Liebermeister, Wolfram and Klipp, Edda},
	title = {Bringing metabolic networks to life: convenience rate law and thermodynamic constraints},
	journal = {Theoretical Biology and Medical Modelling},
	volume = {3},
	pages = {41},
	year = {2006},
	month = {Dec},
	issn = {1742-4682},
	publisher = {BioMed Central},
	doi = {10.1186/1742-4682-3-41}
}

@article{Mason2019Energetic,
	author = {Mason, John C. and Covert, Markus W.},
	title = {An energetic reformulation of kinetic rate laws enables scalable parameter estimation for biochemical networks},
	journal = {Journal of Theoretical Biology},
	volume = {461},
	pages = {145--156},
	year = {2019},
	month = {Jan},
	issn = {0022-5193},
	publisher = {Academic Press},
	doi = {10.1016/j.jtbi.2018.10.041}
}

@article{Gawthrop2014Energy,
	title = {Energy-based analysis of biochemical cycles using bond graphs},
	volume = {470},
	issn = {1364-5021, 1471-2946},
	doi = {10.1098/rspa.2014.0459},
	number = {2171},
	journal = {Proceedings of the Royal Society of London A: Mathematical, Physical and Engineering Sciences},
	author = {Gawthrop, Peter J. and Crampin, Edmund J.},
	month = nov,
	year = {2014},
	pages = {20140459}
}

@article{Gawthrop2020Physically,
	author = {Gawthrop, Peter J. and Cudmore, Peter and Crampin, Edmund J.},
	title = {Physically-plausible modelling of biomolecular systems: A simplified, energy-based model of the mitochondrial electron transport chain},
	journal = {Journal of Theoretical Biology},
	volume = {493},
	pages = {110223},
	year = {2020},
	month = {May},
	issn = {0022-5193},
	publisher = {Academic Press},
	doi = {10.1016/j.jtbi.2020.110223}
}

@book{Schrodinger1962Life,
	author = {Schrödinger, Erwin},
	year = {1962},
	title = {What is life? The physical aspect of the living cell},
	publisher = {The University Press; The Macmillan Company},
	address = {New York},
}

@article{Mallavarapu2008Programming ,
	author = {Mallavarapu, Aneil and Thomson, Matthew and Ullian, Benjamin and Gunawardena, Jeremy},
	title = {Programming with models: modularity and abstraction provide powerful capabilities for systems biology},
	journal = {Journal of The Royal Society Interface},
	volume = {6},
	number = {32},
	pages = {257--270},
	year = {2008},
	month = {Jul},
	publisher = {The Royal Society London},
	doi = {10.1098/rsif.2008.0205}
}

@article{Thomson2009Multistability,
	author = {Thomson, Matthew and Gunawardena, Jeremy},
	title = {Unlimited multistability in multisite phosphorylation systems},
	journal = {Nature},
	volume = {460},
	pages = {274--277},
	year = {2009},
	month = {Jul},
	issn = {1476-4687},
	publisher = {Nature Publishing Group},
	doi = {10.1038/nature08102}
}

@article{Szigeti2018Blueprint,
	author = {Szigeti, Balázs and Roth, Yosef D. and Sekar, John A. P. and Goldberg, Arthur P. and Pochiraju, Saahith C. and Karr, Jonathan R.},
	title = {A blueprint for human whole-cell modeling},
	journal = {Current Opinion in Systems Biology},
	volume = {7},
	pages = {8--15},
	year = {2018},
	month = {Feb},
	issn = {2452-3100},
	publisher = {Elsevier},
	doi = {10.1016/j.coisb.2017.10.005}
}

@article{DelVecchio2013Control,
	author = {Del Vecchio, Domitilla},
	title = {A control theoretic framework for modular analysis and design of biomolecular networks},
	journal = {Annual Reviews in Control},
	volume = {37},
	number = {2},
	pages = {333--345},
	year = {2013},
	month = {Dec},
	issn = {1367-5788},
	publisher = {Pergamon},
	doi = {10.1016/j.arcontrol.2013.09.011}
}

@article{Neal2014Reappraisal,
	author = {Neal, Maxwell L. and Cooling, Michael T. and Smith, Lucian P. and Thompson, Christopher T. and Sauro, Herbert M. and Carlson, Brian E. and Cook, Daniel L. and Gennari, John H.},
	title = {A Reappraisal of How to Build Modular, Reusable Models of Biological Systems},
	journal = {PLOS Computational Biology},
	volume = {10},
	number = {10},
	pages = {e1003849},
	year = {2014},
	month = {Oct},
	issn = {1553-7358},
	publisher = {Public Library of Science},
	doi = {10.1371/journal.pcbi.1003849}
}

@article{Neal2015Semantics,
    author = {Neal, Maxwell L. and Carlson, Brian E. and Thompson, Christopher T. and James, Ryan C. and Kim, Karam G. and Tran, Kenneth and Crampin, Edmund J. and Cook, Daniel L. and Gennari, John H.},
    title = {Semantics-Based Composition of Integrated Cardiomyocyte Models Motivated by Real-World Use Cases},
    journal = {PLOS ONE},
    volume = {10},
    number = {12},
    pages = {e0145621},
    year = {2015},
    month = {Dec},
    issn = {1932-6203},
    publisher = {Public Library of Science},
    doi = {10.1371/journal.pone.0145621}
}

@article{Lopez2013Programming,
	author = {Lopez, Carlos F. and Muhlich, Jeremy L. and Bachman, John A. and Sorger, Peter K.},
	title = {Programming biological models in {Python} using {PySB}},
	journal = {Molecular Systems Biology},
	volume = {9},
	number = {1},
	pages = {646},
	year = {2013},
	month = {Jan},
	issn = {1744-4292},
	publisher = {John Wiley {\&} Sons, Ltd},
	doi = {10.1038/msb.2013.1}
}

@article{Daly2018Reproducible,
	author = {Daly, Aidan C. and Clerx, Michael and Beattie, Kylie A. and Cooper, Jonathan and Gavaghan, David J. and Mirams, Gary R.},
	title = {Reproducible model development in the cardiac electrophysiology {Web Lab}},
	journal = {Progress in Biophysics and Molecular Biology},
	volume = {139},
	pages = {3--14},
	year = {2018},
	month = {Nov},
	issn = {0079-6107},
	publisher = {Pergamon},
	doi = {10.1016/j.pbiomolbio.2018.05.011}
}

@article{Gawthrop2017Chemiosmotic,
	author = {Gawthrop, Peter J.},
	title = {Bond Graph Modeling of Chemiosmotic Biomolecular Energy Transduction},
	journal = {IEEE Transactions on NanoBioscience},
	volume = {16},
	number = {3},
	pages = {177--188},
	year = {2017},
	month = {Feb},
	publisher = {IEEE},
	doi = {10.1109/TNB.2017.2674683}
}

@article{Gawthrop2020Bioelectricity,
	author = {Gawthrop, Peter J. and Pan, Michael},
	title = {Network Thermodynamical Modeling of Bioelectrical Systems: A Bond Graph Approach},
	journal = {Bioelectricity},
	year = {2020},
	month = {Dec},
	issn = {2576-3105},
	publisher = {Mary Ann Liebert, Inc., publishers},
	doi = {10.1089/bioe.2020.0042}
}

@article{Tran2015Regulation,
	author = {Tran, Kenneth and Loiselle, Denis S. and Crampin, Edmund J.},
	title = {Regulation of cardiac cellular bioenergetics: mechanisms and consequences},
	journal = {Physiological Reports},
	volume = {3},
	number = {7},
	pages = {e12464},
	year = {2015},
	month = {Jul},
	issn = {2051-817X},
	publisher = {John Wiley {\&} Sons, Ltd},
	doi = {10.14814/phy2.12464}
}

@incollection{Crampin2020Energetics,
    author = {Crampin, Edmund J and Pan, Michael and Tran, Kenneth and Ghosh, Shourya and Rajagopal, Vijay},
    title = {Modelling cardiomyocyte energetics},
    booktitle = {Modeling and Simulating Cardiac Electrical Activity},
    editor = {Krogh-Madsen, Trine and Christini, David J},
    publisher = {IOP Publishing},
    year = {2020},
    series = {2053-2563},
    type = {Book Chapter},
    pages = {6-1 to 6-45},
    doi = {10.1088/978-0-7503-2064-1ch6},
    isbn = {978-0-7503-2064-1}
}

@article{Chylek2014Rule,
	author = {Chylek, Lily A. and Harris, Leonard A. and Tung, Chang-Shung and Faeder, James R. and Lopez, Carlos F. and Hlavacek, William S.},
	title = {Rule-based modeling: a computational approach for studying biomolecular site dynamics in cell signaling systems},
	journal = {WIREs Syst. Biol. Med.},
	volume = {6},
	number = {1},
	pages = {13--36},
	year = {2014},
	month = {Jan},
	issn = {1939-5094},
	publisher = {John Wiley {\&} Sons, Ltd},
	doi = {10.1002/wsbm.1245}
}

@book{Paynter1961a,
  title={Analysis and design of engineering systems},
  author={Paynter, Henry M},
  year={1961},
  publisher={MIT. Press},
  address={Cambridge, Mass.}
}

@article{Gawthrop2007aa,
	title = {Bond-graph modeling},
	volume = {27},
	doi = {10.1109/MCS.2007.338279},
	number = {2},
	journal = {IEEE Control Systems Magazine},
	author = {Gawthrop, P.J. and Bevan, G.P.},
	year = {2007},
	pages = {24--45}
}

@book{KarMarRos12,
	author = {Karnopp, Dean C. and Margolis, Donald L. and Rosenberg, Ronald C.},
	title = {System Dynamics: Modeling, Simulation, and Control of
		Mechatronic Systems},
	journal = {Wiley Online Library},
	year = {2012},
	month = {Feb},
	isbn = {978-0-47088908-4},
	doi = {10.1002/9781118152812},
    publisher={John Wiley \& Sons},
    edition={5th},
	address={Hoboken, New Jersey}
}

@article{Rac2017,
	author = {Rackauckas, Christopher and Nie, Qing},
	title = {DifferentialEquations.jl {\textendash} A Performant and Feature-Rich Ecosystem for Solving Differential Equations in Julia},
	journal = {Journal of Open Research Software},
	volume = {5},
	number = {1},
    pages={15},
	year = {2017},
	month = {May},
	issn = {2049-9647},
	publisher = {Ubiquity Press},
	doi = {10.5334/jors.151}
}

@article{sundials,
	author = {Hindmarsh, Alan C. and Brown, Peter N. and Grant, Keith E. and Lee, Steven L. and Serban, Radu and Shumaker, Dan E. and Woodward, Carol S.},
	title = {{SUNDIALS}: Suite of nonlinear and differential/algebraic equation solvers},
	journal = {ACM Transactions on Mathematical Software},
	volume = {31},
	number = {3},
	pages = {363--396},
	year = {2005},
	month = {Sep},
	issn = {0098-3500},
	publisher = {Association for Computing Machinery},
	doi = {10.1145/1089014.1089020}
}

@article{Julia,
	author = {Bezanson, Jeff and Edelman, Alan and Karpinski, Stefan and Shah, Viral B.},
	title = {Julia: A Fresh Approach to Numerical Computing},
	journal = {SIAM Review},
	year = {2017},
    volume={59},
    number={1},
    pages={65–-98},
	month = {Feb},
	publisher = {Society for Industrial and Applied Mathematics},
	url = {https://epubs.siam.org/doi/abs/10.1137/141000671}
}

@article{Pan2018aa,
	author = {Pan, Michael and Gawthrop, Peter J. and Tran, Kenneth and Cursons, Joseph and Crampin, Edmund J.},
	title = {A thermodynamic framework for modelling membrane transporters},
	journal = {Journal of Theoretical Biology},
	volume = {481},
	pages = {10--23},
	year = {2019},
	month = {Nov},
	issn = {0022-5193},
	publisher = {Academic Press},
	doi = {10.1016/j.jtbi.2018.09.034}
}

@article{Pan2018ab,
	author = {Pan, Michael and Gawthrop, Peter J. and Tran, Kenneth and Cursons, Joseph and Crampin, Edmund J.},
	title = {Bond graph modelling of the cardiac action potential: implications for drift and non-unique steady states},
	journal = {Proceedings of the Royal Society A: Mathematical, Physical and Engineering Sciences},
	year = {2018},
	volume = {474},
	number = {2214},
	pages = {20180106},
	month = {Jun},
	publisher = {The Royal Society Publishing},
	doi = {10.1098/rspa.2018.0106}
}

@article{Dan2018,
	author = {Hurley, Daniel G. and Cursons, Joseph and Faria, Matthew and Budden, David M. and Rajagopal, Vijay and Crampin, Edmund J.},
	title = {Reference environments: A universal tool for reproducibility in computational biology},
	journal = {arXiv},
	year = {2018},
	month = {Oct},
	eprint = {1810.03766},
	url = {https://arxiv.org/abs/1810.03766v1}
}

@article{Perez2007,
	author = {Perez, Fernando and Granger, Brian E.},
	title = {{IPython}: A System for Interactive Scientific Computing},
	journal = {Computing in Science {\&} Engineering},
	volume = {9},
	number = {3},
	pages = {21--29},
	year = {2007},
	month = {May},
	issn = {1558-366X},
	doi = {10.1109/MCSE.2007.53}
}

@article{Cudmore2015Feb,
	author = {Cudmore, P. and Holmes, C. A.},
	title = {Phase and amplitude dynamics of nonlinearly coupled oscillators},
	journal = {Chaos: An Interdisciplinary Journal of Nonlinear Science},
	volume = {25},
	number = {2},
	pages = {023110},
	year = {2015},
	month = {Feb},
	issn = {1054-1500},
	publisher = {American Institute of Physics},
	doi = {10.1063/1.4908604}
}

@article{Diaz2011Multiscale,
	author = {Díaz-Zuccarini, Vanessa and Pichardo-Almarza, César},
	title = {On the formalization of multi-scale and multi-science processes for integrative biology},
	journal = {Interface Focus},
	volume = {1},
	number = {3},
	pages = {426--437},
	year = {2011},
	month = {Jun},
	issn = {2042-8901},
	publisher = {The Royal Society},
	doi = {10.1098/rsfs.2010.0038}
}

@article{deBono2018Multiscale,
	author = {de Bono, B. and Safaei, S. and Grenon, P. and Hunter, P.},
	title = {Meeting the multiscale challenge: representing physiology processes over ApiNATOMY circuits using bond graphs},
	journal = {Interface Focus},
	volume = {8},
	number = {1},
	pages = {20170026},
	year = {2018},
	month = {Feb},
	publisher = {The Royal Society},
	doi = {10.1098/rsfs.2017.0026}
}

@article{Gawthrop2015Hierarchical,
	author = {Gawthrop, Peter J. and Cursons, Joseph and Crampin, Edmund J.},
	title = {Hierarchical bond graph modelling of biochemical networks},
	journal = {Proceedings of the Royal Society A: Mathematical, Physical and Engineering Sciences},
	year = {2015},
	volume = {471},
	number = {2184},
	pages = {20150642},
	month = {Dec},
	publisher = {The Royal Society Publishing},
	doi = {10.1098/rspa.2015.0642}
}

@article{hurley_virtual_2015,
	title = {Virtual reference environments: a simple way to make research reproducible},
	volume = {16},
	number = {5},
	journal = {Briefings in Bioinformatics},
	author = {Hurley, Daniel G and Budden, David M and Crampin, Edmund J},
	month = sep,
	year = {2015},
	pages = {901--903}
}

@article{Kohl2010Systems,
	author = {Kohl, P. and Crampin, E. J. and Quinn, T. A. and Noble, D.},
	title = {Systems Biology: An Approach},
	journal = {Clinical Pharmacology {\&} Therapeutics},
	volume = {88},
	number = {1},
	pages = {25--33},
	year = {2010},
	month = {Jul},
	issn = {0009-9236},
	publisher = {John Wiley {\&} Sons, Ltd},
	doi = {10.1038/clpt.2010.92}
}

@article{Karr2012Whole,
	author = {Karr, Jonathan R. and Sanghvi, Jayodita C. and Macklin, Derek N. and Gutschow, Miriam V. and Jacobs, Jared M. and Bolival, Benjamin and Assad-Garcia, Nacyra and Glass, John I. and Covert, Markus W.},
	title = {A Whole-Cell Computational Model Predicts Phenotype from Genotype},
	journal = {Cell},
	volume = {150},
	number = {2},
	pages = {389--401},
	year = {2012},
	month = {Jul},
	issn = {0092-8674},
	publisher = {Elsevier},
	doi = {10.1016/j.cell.2012.05.044}
}

@article{Macklin2020Simultaneous,
	author = {Macklin, Derek N. and Ahn-Horst, Travis A. and Choi, Heejo and Ruggero, Nicholas A. and Carrera, Javier and Mason, John C. and Sun, Gwanggyu and Agmon, Eran and DeFelice, Mialy M. and Maayan, Inbal and Lane, Keara and Spangler, Ryan K. and Gillies, Taryn E. and Paull, Morgan L. and Akhter, Sajia and Bray, Samuel R. and Weaver, Daniel S. and Keseler, Ingrid M. and Karp, Peter D. and Morrison, Jerry H. and Covert, Markus W.},
	title = {Simultaneous cross-evaluation of heterogeneous E. coli datasets via mechanistic simulation},
	journal = {Science},
	volume = {369},
	number = {6502},
	pages = {eaav3751},
	year = {2020},
	month = {Jul},
	issn = {0036-8075},
	publisher = {American Association for the Advancement of Science},
	doi = {10.1126/science.aav3751}
}

@article{Noble2001Cardiac,
	author = {Noble, D. and Rudy, Y.},
	title = {Models of cardiac ventricular action potentials: iterative interaction between experiment and simulation},
	journal = {Philosophical Transactions of the Royal Society of London. Series A: Mathematical, Physical and Engineering Sciences},
	volume = {359},
	number = {1783},
	pages = {1127--1142},
	year = {2001},
	month = {Jun},
	issn = {1471-2962},
	publisher = {The Royal Society},
	doi = {10.1098/rsta.2001.0820}
}

@article{Hund2001Ionic,
	author = {Hund, Thomas J. and Kucera, Jan P. and Otani, Niels F. and Rudy, Yoram},
	title = {Ionic Charge Conservation and Long-Term Steady State in the Luo{\textendash}Rudy Dynamic Cell Model},
	journal = {Biophysical Journal},
	volume = {81},
	number = {6},
	pages = {3324--3331},
	year = {2001},
	month = {Dec},
	issn = {0006-3495},
	publisher = {Cell Press},
	doi = {10.1016/S0006-3495(01)75965-6}
}

@article{Safaei2016Circulation,
	author = {Safaei, Soroush and Bradley, Christopher P. and Suresh, Vinod and Mithraratne, Kumar and Muller, Alexandre and Ho, Harvey and Ladd, David and Hellevik, Leif R. and Omholt, Stig W. and Chase, J. Geoffrey and Müller, Lucas O. and Watanabe, Sansuke M. and Blanco, Pablo J. and de Bono, Bernard and Hunter, Peter J.},
	title = {Roadmap for cardiovascular circulation model},
	journal = {Journal of Physiology},
	volume = {594},
	number = {23},
	pages = {6909--6928},
	year = {2016},
	month = {Dec},
	issn = {0022-3751},
	publisher = {John Wiley {\&} Sons, Ltd},
	doi = {10.1113/JP272660}
}

@article{Saa2015Sampling,
	author = {Saa, Pedro and Nielsen, Lars K.},
	title = {A General Framework for Thermodynamically Consistent Parameterization and Efficient Sampling of Enzymatic Reactions},
	journal = {PLOS Computational Biology},
	volume = {11},
	number = {4},
	pages = {e1004195},
	year = {2015},
	month = {Apr},
	issn = {1553-7358},
	publisher = {Public Library of Science},
	doi = {10.1371/journal.pcbi.1004195}
}


@article{crampin_multi-scale_2004,
	title = {Multi-scale modelling and the {IUPS} physiome project},
	volume = {35},
	number = {7},
	journal = {Journal of Molecular Histology},
	author = {Crampin, Edmund J and Smith, Nicolas P and Hunter, Peter J},
	year = {2004},
	pages = {707--714}
}

@article{hunter_virtual_2016,
	title = {{The Virtual Physiological Human}: {The Physiome Project} Aims to Develop Reproducible, Multiscale Models for Clinical Practice},
	volume = {7},
	issn = {2154-2287},
	shorttitle = {The {Virtual} {Physiological} {Human}},
	doi = {10.1109/MPUL.2016.2563841},
	number = {4},
	urldate = {2021-03-12},
	journal = {IEEE Pulse},
	author = {Hunter, Peter},
	month = jul,
	year = {2016},
	pages = {36--42}
}

@article{Shahidi2021Semantic,
	author = {Shahidi, Niloofar and Pan, Michael and Safaei, Soroush and Tran, Kenneth and Crampin, Edmund J. and Nickerson, David P.},
	title = {Hierarchical semantic composition of biosimulation models using bond graphs},
	journal = {bioRxiv},
	pages = {2021.03.09.434672},
	year = {2021},
	month = {Mar},
	publisher = {Cold Spring Harbor Laboratory},
	eprint = {2021.03.09.434672},
	doi = {10.1101/2021.03.09.434672},
}
