## Supplementary material for "Analysing and simulating energy-based models in biology using BondGraphTools": LaTeX source for manuscript: BondGraphTools.tex

%%%%%%%%%%%%%%%%%%%%%%% file template.tex %%%%%%%%%%%%%%%%%%%%%%%%%
%
% This is a  template file for the LaTeX package SVJour3 width change file svepjc3.clo
% for Springer journal:
% The European Physical Journal E
%
% Copy it to a new file with a new name and use it as the basis
% for your article. Delete % signs as needed.
%
% This template includes a few options for different layouts and
% content for various journals. Please consult a previous issue of
% your journal as needed.
%
%%%%%%%%%%%%%%%%%%%%%%%%%%%%%%%%%%%%%%%%%%%%%%%%%%%%%%%%%%%%%%%%%%%
%
%
\RequirePackage{fix-cm}
%
\documentclass[twocolumn,epjc3]{svjour3}  
%
\smartqed  % flush right qed marks, e.g. at end of proof
%
\RequirePackage{graphicx}
\usepackage{amsmath}
\usepackage{amsfonts}
\usepackage{xcolor}
\usepackage{siunitx}
\usepackage{booktabs}
\usepackage{multirow}
\usepackage{microtype}
\usepackage{hyperref}
\usepackage{listings}
\usepackage{cite}
\usepackage[switch,mathlines,pagewise]{lineno}
\definecolor{codebg}{HTML}{F6F6F6}
\definecolor{codeframe}{HTML}{CCCCCC}
\lstset{
	language=Python,
	backgroundcolor=\color{codebg},
	frame=single,
	keywordstyle=\bfseries\color{green!40!black},
	basicstyle=\scriptsize\ttfamily,
	stringstyle=\color{orange},
	rulecolor=\color{black},  
    keepspaces=true,
    tabsize=4    
}

%\linenumbers
%\renewcommand{\linenumberfont}{\sffamily\tiny\color{gray}}
%\let\oldalign\align
%\let\oldendalign\endalign
%\renewenvironment{align}
%  {\linenomathNonumbers\oldalign}
%  {\oldendalign\endlinenomath}

% Math Macros
\newcommand{\Epsilon}{\mathcal{E}}
\newcommand{\Chi}{\mathcal{X}}
\newcommand{\D}[2]{\frac{\mathrm{d} #1}{\mathrm{d} #2}}
\newcommand{\DD}[2]{\frac{\mathrm{d}^2 #1}{\mathrm{d} #2^2}}
\newcommand{\DDT}{\frac{\mathrm{d}}{\mathrm{d} t}}
\newcommand{\df}[1]{\mspace{2mu}  \mathrm{d}#1}
\renewcommand{\P}[2]{\frac{\partial #1}{\partial #2}}
\newcommand{\PP}[2]{\frac{\partial^2 #1}{\partial #2 ^2}}
\newcommand{\e}{\mathrm{e}}
\newcommand{\bld}[1]{\mathbf{#1}}
\newcommand{\lie}[1]{\mathcal{L}_{#1}}
\newcommand{\dW}[1]{\circ\mathrm{dW}_{#1}}
\newcommand{\Exp}{\mathbf{Exp}}
\newcommand{\Ln}{\mathbf{Ln}}
\newcommand{\reals}{\mathbb{R}}
\newcommand{\I}{\mathrm{i}}
\newcommand{\BGT}{\texttt{BondGraphTools}}
\newcommand{\Ce}{\textbf{Ce}}
\newcommand{\Se}{\textbf{Se}}
\newcommand{\zero}{\textbf{0}}
\newcommand{\one}{\textbf{1}}

%
% \RequirePackage{mathptmx}      % use Times fonts if available on your TeX system
%
% insert here the call for the packages your document requires
%\RequirePackage{latexsym}
%\RequirePackage[numbers,sort&compress]{natbib}
%\RequirePackage[colorlinks,citecolor=blue,urlcolor=blue,linkcolor=blue]{hyperref}
% etc.
%
% please place your own definitions here and don't use \def but
% \newcommand{}{}
%
\journalname{Preprint}
%
\begin{document}

\title{Analysing and simulating energy-based models in biology using BondGraphTools}
\subtitle{}

%\titlerunning{Short form of title}        % if too long for running head

\author{Peter Cudmore\thanksref{addr1,addr2}
            \and
        Michael Pan\thanksref{addr1,addr2}
          \and
        Peter J. Gawthrop\thanksref{addr1} 
         \and
        \mbox{Edmund J. Crampin}\thanksref{addr1,addr2,addr3,e1}
}


%\thankstext{t1}{Grants or other notes
%about the article that should go on the front page should be
%placed here. General acknowledgments should be placed at the end of the article.

%\authorrunning{Short form of author list} % if too long for running head

\institute{Systems Biology Laboratory, School of Mathematics and Statistics, and Department of Biomedical Engineering, University of Melbourne, Parkville, Victoria 3010 \label{addr1}
           \and
           ARC Centre of Excellence in Convergent Bio-Nano Science and Technology, Faculty of Engineering and Information Technology, University of Melbourne, Parkville, Victoria 3010 \label{addr2}
           \and
           School of Medicine, Faculty of Medicine, Dentistry and Health Sciences, University of Melbourne, Parkville, Victoria 3010 \label{addr3}
}

\date{Received: date / Accepted: date}
% The correct dates will be entered by the editor

\abstractdc{Like all physical systems, biological systems are constrained by the laws of physics. However, mathematical models of biochemistry frequently neglect the conservation of energy, leading to unrealistic behaviour. Energy-based models that are consistent with conservation of mass, charge and energy have the potential to aid the understanding of complex interactions between biological components, and are becoming easier to develop with recent advances in experimental measurements and databases. In this paper, we motivate the use of bond graphs (a modelling tool from engineering) for energy-based modelling and introduce, \BGT, a Python library for constructing and analysing bond graph models. We use examples from biochemistry to illustrate how \BGT\ can be used to automate model construction in systems biology while maintaining consistency with the laws of physics.}

\maketitle

%
\section{Introduction}
Biological systems are highly complex, composed of numerous interacting components. Systems biology aims to explain how complex phenomena arise from interactions between these components \cite{Kohl2010Systems}. While mathematical models have been long been used to understand biology, efforts to integrate these models together have gathered momentum since the turn of the century, together with the availability of omics data. Examples of this approach include whole-cell models that simulate the evolution of each biomolecule through time, \cite{Karr2012Whole,Szigeti2018Blueprint,Macklin2020Simultaneous} through to multiscale physiological models \cite{crampin_multi-scale_2004,hunter_virtual_2016}.

Like any physical system, interactions between biological components are governed by the laws of physics; notably conservation of mass, charge and energy. Many existing approaches in the field of physiology take advantage of such conservation laws as a means of coupling models together. This has been used to great success in using conservation of mass and charge to couple models of electrophysiology \cite{Noble2001Cardiac,Hund2001Ionic} and coupling boundary conditions in models of fluid flow \cite{Safaei2016Circulation}.

Energy is fundamental to biological systems, as it is to physical systems. In his book \textit{What is Life?}, the physicist Erwin Schrodinger wrote that living systems are in a constant fight against a decay to thermodynamic equilibrium, and must feed on free energy from the environment to avoid this eventual approach to equilibrium\cite{Schrodinger1962Life}. \textit{Energy-based models} that account for the conservation of mass, charge and energy have the potential to gain a deeper understanding of the forces driving biological networks. However, in the field of systems biology, energy conservation is commonly ignored, and many models are thermodynamically inconsistent. As discussed by Soh and Hatzimanikatis \cite{Soh2010Network}, difficulties in measuring the thermodynamic parameters have slowed the uptake of energy-based models. However, recent developments in thermodynamic databases such as eQuilibrator \cite{Flamholz2012Jan} have made constructing thermodynamically detailed models more tractable. 

In addition to developments in data availability, there have also been recent advancements in methodologies for constructing energy-based models. The approach was first introduced to biology by Oster \textit{et al.} \cite{Oster1973Network}, but more recent examples include application to systems biology by Ederer and Gilles \cite{Ederer2007Thermodynamically}, Liebermeister and Klipp \cite{Liebermeister2006Bringing} and Gawthrop and Crampin \cite{Gawthrop2014Energy}. Whereas traditional kinetic models in biochemistry depend on complex detailed balance constraints to account for energy conservation, a key advantage of energy-based models is that individual parameters can be individually perturbed without violating thermodynamic consistency. Such developments are starting to find application in whole-cell modelling, with Mason and Covert \cite{Mason2019Energetic} finding that parameterising models in a thermodynamically consistent fashion can in fact lead to more robust parameter identification. 

The development and analysis of energy-based models can be significantly simplified if one knows the energetic properties of each component in a system and how the components are connected. To this effect, bond graphs have proved to be valuable tools for enabling the energy-based approach. Bond graphs were invented by Henry Paynter as a method of modelling multi-physics systems in engineering. They were subsequently proposed as a modelling approach in biology by Oster \textit{et al.} \cite{Oster1973Network}, and more recently developed by Gawthrop and Crampin \cite{Gawthrop2014Energy}. Because bond graphs are based on a graphical representation, they allow the complexities of energy-based modelling to be abstracted away so that a modeller need only deal with a map (graph) of biological components and their connections.

However, despite recent developments in bond graph theory, software tools for constructing and simulating bond graph models, in particular of biological systems, have not kept up with recent developments in software practices. Almost all of the existing software implementations of bond graph modelling are graphical in nature and within a proprietary or isolated software ecosystem. For some applications, this is beneficial; vendors can provide a standardised visual interface with integrated analysis tools. However, for systems biology, the existing software lacks the capacity for automation, is hampered by restricted access to source code and is challenging to integrate with existing algorithms. Hence, there is need for a symbolic modelling toolkit that is open-source, easy to integrate into existing workflows, and written in an accessible and widely used scripting language. In particular, there is a need for infrastructure to support automated model building and simplification.

Here we introduce \BGT, a python library for building and manipulating symbolic models of complex physical systems, built upon the standard scientific python libraries.
The \BGT\ package is different from existing software in both design and implementation in that it:
\begin{enumerate}
    \item is explicitly based on physical modelling principles;
    \item provides an \emph{application program interface} (API) as opposed to a graphical user interface (GUI);
    \item is designed for \emph{symbolic} model composition and order reduction, as opposed to being primarily a numerical toolkit;
    \item is intended to be used in conjunction with the standard python libraries, as opposed to being used as a stand-alone software package;
    \item allows for modification and integration as it is open source, version controlled, and readily available, instead of closed-source and proprietary;
    \item is designed with modularity and extensibility in mind, as opposed to being a monolithic software suite.
\end{enumerate}

The aims of this paper are threefold: firstly to use some simple examples to motivate the use of an energy-based approach for modelling cellular biochemistry (Sect. \ref{sec:thermodynamic_cycles}); secondly to illustrate how bond graphs facilitate this approach (Sect. \ref{sec:bg_modelling}); and thirdly to introduce the \BGT\ package for simulating energy-based models in systems biology (Sects. \ref{sec:bgt}--\ref{sec:reductions}). We apply this energy-based approach to a model of multisite phosphorylation to illustrate the advantages of this approach for large-scale modelling (Sect. \ref{sec:example}).

\section{A primer on modelling biochemical cycles} \label{sec:thermodynamic_cycles}
Biochemical networks contain numerous thermodynamic cycles. Here we motivate the need to consider thermodynamics when such cycles exist. As a simple example, consider the reactions
\begin{align}
	\mathrm{X \rightleftharpoons Y \rightleftharpoons Z \rightleftharpoons X}
\end{align}
Assuming the law of mass action, the system can be described by the differential equations
\begin{subequations}
\begin{align}
    \dot{x}_X &= v_3 - v_1 \\
    \dot{x}_Y &= v_1 - v_2 \\
    \dot{x}_Z &= v_2 - v_3 
\end{align} \label{eq:mass_conservation}
\end{subequations}
where $x_s$ is the concentration of species $s \in \{\mathrm{X,Y,Z}\}$. The reaction fluxes are defined as
\begin{subequations}
\begin{align}
    v_1 &= k_1^+ x_X - k_1^- x_Y \\
    v_2 &= k_2^+ x_Y - k_2^- x_Z \\
    v_3 &= k_3^+ x_Z - k_3^- x_X 
\end{align} \label{eq:reaction_kinetics}
\end{subequations}
where $k_i^\pm$ are kinetic constants.

It is clear that at steady state, $v_1 = v_2 = v_3$. However, because the system is closed, this steady state must satisfy the stricter constraint that $v_1 = v_2 = v_3 = 0$. Otherwise, the reactions would dissipate heat without any energy being input into the system, violating the second law of thermodynamics. With this in mind, from Eq. \ref{eq:reaction_kinetics},
\begin{subequations}
\begin{align}
    k_1^+ x_X &= k_1^- x_Y \\
    k_2^+ x_Y &= k_2^- x_Z \\
    k_3^+ x_Z &= k_3^- x_X
\end{align}
\end{subequations}
By taking the product of the three equations, it is easy to see that
\begin{align}
    k_1^+ x_X k_2^+ x_Y k_3^+ x_Z = k_1^- x_Y k_2^- x_Z k_3^- x_X
    \Rightarrow \frac{k_1^+ k_2^+ k_3^+}{k_1^- k_2^- k_3^-} = 1 \label{eq:detailed_balance}
\end{align}
Since Eq. \ref{eq:detailed_balance} is independent of the concentrations of species, it is a constraint on the kinetic parameters. In other words, the parameters $k_i^\pm$ cannot be independently chosen and must satisfy Eq. \ref{eq:detailed_balance} to avoid describing a system analogous to a perpetual motion machine \cite{Ederer2007Thermodynamically}. This poses a few issues:
\begin{enumerate}
    \item Detailed balance constraints become substantially more difficult to calculate for complex biochemical networks \cite{Ederer2007Thermodynamically}
    \item It is challenging to specify kinetic parameters (or infer from data) while adhering to detailed balance constraints \cite{Saa2015Sampling}
\end{enumerate}

An alternative method of representing biochemical kinetics is the energy-based approach, which explicitly accounts for the fact that reaction fluxes arise from chemical potentials \cite{Gawthrop2014Energy}. Here, each of the species has an associated chemical potential
\begin{subequations}
\begin{align}
    \mu_X &= RT \ln (K_X x_X) \\
    \mu_Y &= RT \ln (K_Y x_Y) \\
    \mu_Z &= RT \ln (K_Z x_Z) 
\end{align} \label{eq:XYZ_mu}
\end{subequations}
%
where $R = 8.314\ \si{J/mol}$ is the ideal gas constant, $T$ [K] is the temperature and $K_i$ are the thermodynamic constants of the species. The chemical potentials are related to reaction rates through the Marcelin-de Donder equation
\begin{subequations}
\begin{align} 
    v_1 &= r_1 (e^{\mu_X/RT} - e^{\mu_Y/RT}) \\
    v_2 &= r_2 (e^{\mu_Y/RT} - e^{\mu_Z/RT}) \\
    v_3 &= r_3 (e^{\mu_Z/RT} - e^{\mu_X/RT}) 
\end{align} \label{eq:XYZ_v}
\end{subequations}
where $r_i$ [$\si{mol/s}$] is the rate parameter for reaction $i$. By substituting Eq. \ref{eq:XYZ_mu} into Eq. \ref{eq:XYZ_v}, the equations can be recast in the same form as Eq. \ref{eq:mass_conservation}, but with the reaction rates
\begin{subequations}
\begin{align}
    v_1 &= r_1 K_X x_X - r_1 K_Y x_Y \label{eq:v1}\\
    v_2 &= r_2 K_Y x_Y - r_2 K_Z x_Z \\
    v_3 &= r_3 K_Z x_Z - r_3 K_X x_X
\end{align} \label{eq:flux_energetic}
\end{subequations}
While these equations exhibit the same mass-action behaviour as the kinetic approach, the thermodynamic approach is immune to thermodynamic inconsistencies; by comparing coefficients between Eqs. \ref{eq:reaction_kinetics} and \ref{eq:flux_energetic},
\begin{align}
    \frac{k_1^+ k_2^+ k_3^+}{k_1^- k_2^- k_3^-} = \frac{r_1 K_X r_2 K_Y r_3 K_Z}{r_1 K_Y r_2 K_Z r_3 K_X} = 1
\end{align}
that is, the parameters in the energetic formulation automatically satisfy the detailed balance constraint in Eq. \ref{eq:detailed_balance} and thus the parameters are free to be specified independently. This has been found to aid in estimating the parameters for a model of glycolysis \cite{Mason2019Energetic}.

While the above example considered an closed system, biological systems in general continuously dissipate energy. The equilibrium constraint is broken when external chemical species are allowed to interact with the cycle. For example, we now consider the alternative reaction network
\begin{subequations}
\begin{align}
    \mathrm{X + A} &\rightleftharpoons \mathrm{Y} \\
    \mathrm{Y} &\rightleftharpoons \mathrm{Z} \\
    \mathrm{Z} &\rightleftharpoons \mathrm{X + B}
\end{align}
\end{subequations}
and assume that $\mathrm{A}$ and $\mathrm{B}$ are kept at constant concentrations through external flows. This is a biochemical cycle frequently used by enzymes. The reaction rates are 
\begin{subequations}
\begin{align}
    v_1 &= r_1 K_X x_X K_A \bar{x}_A - r_1 K_Y x_Y \\
    v_2 &= r_2 K_Y x_Y - r_2 K_Z x_Z\\
    v_3 &= r_3 K_Z x_Z - r_3 K_X x_X K_B \bar{x}_B
\end{align}  \label{eq:open_v}
\end{subequations}
where $K_A$ and $K_B$ are the species constants for A and B respectively and $\bar{x}_A$ and $\bar{x}_B$ are the constant concentrations of A and B.
By comparing coefficients with Eq. \ref{eq:reaction_kinetics},
\begin{align}
    \frac{k_1^+ k_2^+ k_3^+}{k_1^- k_2^- k_3^-} = \frac{r_1 K_1 K_A \bar{x}_A r_2 K_2 r_3 K_3}{r_1 K_2 r_2 K_3 r_3 K_1 K_B \bar{x}_B} = \frac{K_A \bar{x}_A}{K_B \bar{x}_B}
\end{align}
which is not 1 in general, and therefore admits a non-equilibrium steady state. Thus the addition of external species provides a source of energy to the system, allowing the system to dissipate energy under steady state conditions.

\section{Introduction to bond graph modelling}
\label{sec:bg_modelling}
%\subsection{Introduction to bond graphs}
Bond graphs provide an abstracted and network-based framework for understanding and developing energy-based models. The main principle of bond graph modelling is that the laws of physics  provide a justifiable interface between different models of physical subsystems. In particular any \emph{connection} between two models of physical processes
%, that is any non-physical feature of the modelling framework set up to allow disparate sub-models to interface, 
must conserve energy. 

Bond graphs are a port-based modelling approach that describes the flow of power through a network of energy storage and dissipation sub-systems (such as species and reactions respectively). Ports attached to a particular subsystem can be related via an energy conserving `power bond' and are defined in terms of \emph{force-like} 
%efforts (or 
potentials $e$ (also known in the bond graph literature as `efforts'), and \emph{flux-like} flows $f$ such that the (signed) power transfer at any instant is $P=ef$. In bond graphs, variables such as voltage, pressure and chemical potential are potentials; while current, mass flow and molar flow rate are flow variables. 
Figure~\ref{fig:power_bond} shows an example of a power bond. Here, the power entering system A is given by $P_{\text{A};j} = e_jf_j$, similarly for B; $P_{\text{B};k} = e_kf_k$, such that power is conserved between systems, \textit{i.e.} $P_{\text{A};j}+P_{\text{B};k} = 0$. 
The directed power transfer through the bond is therefore 
\begin{align}
P_{bond}=ef,\ \text{with}\ e = e_j = e_k \ \text{and}\ f = -f_j = f_k.
\label{eq:bond}
\end{align}
The internal behaviour of systems A and B are captured via the constitutive relationships $\Phi_\text{A}$ and $\Phi_\text{B}$, which store or dissipate energy.
We refer interested readers to the tutorial by Gawthrop and Bevan~\cite{Gawthrop2007aa} for an overview of bond graph modelling and for an in depth treatment see~\cite{KarMarRos12}.
%
%One interpretation of bond graphs is as formalising modelling via analogy. In bond graphs, variables such as voltage, pressure and chemical potential are potentials, while current, mass flow and molar flow rate are flow variables. This allows insights from formal disciplines such as electrical engineering to be applied to biological systems.
\begin{figure}
	\centering
	\includegraphics{figure1.pdf}
	\caption{The transfer of power through a bond between port $j$ on system A and port $k$ on system B.
		\label{fig:power_bond}}
\end{figure}
%More recently, there has been an effort spearheaded by van der Schaft~\cite{VanderSchaft2014b} and others, to integrate port-based bond graphs approaches with modern geometric mechanics resulting in theory of port-Hamiltonian systems. This follows from the observation that for conservative systems effort and flow variables can be related to the time derivatives of an energy storage function. 

In order to model complex systems, we must first have an idea of what the model subsystems are, and how they should be connected.
All bond graph models have:
\begin{itemize}
	\item a number of internal components,
	\item a (possibly zero) number of external power \emph{ports} each of which has two associated variables, a potential-like $e$ (voltage, force, chemical potential, etc.) and a flow-like $f$ (current, velocity, molar flow rate, etc.) such that power $P=ef$ is positive when the process is consuming or accumulating energy through that particular port,
	\item a set of constitutive relations that characterise the behaviour for a particular model and are either derived from internal components or specified a-priori; for example generalised linear dissipation (friction, Ohm's law) which relates the effort and flow via the implicit relation $0 = e - Rf$. We note that the power entering this model $P = ef = Rf^2$ is positive semi-definite for $R>0$ indicating, as one would expect, that resistance always consumes power. 
	\item a (possibly zero) number of parameters/controls which govern the behaviour of that particular model. One would consider $R$ in the above example to be a parameter if $R$ is constant, and a control in all other cases.
	\item a (possibly zero) number of state variables (with associated derivatives), related to power ports via the model constitutive relations. For example, a linear potential energy storage has governing equations 
	$Ce - x = 0,\qquad f - \dot{x} =0,\quad \implies P = \frac{1}{C}x\dot{x} $
	so that the energy $E(t)$ stored in the state variable $x(t)$ at a particular time $t$ in that component (up to a constant) is $E(t) =\int_{0}^tP\df{t} = x(t)^2/(2C)$. 
\end{itemize}

Thus building a bond graph model consists of instantiating the models one wishes to use, defining parent-child relationships, then specifying energy sharing between ports. The resulting equations can then be manipulated and simplified to give rise to a set of differential equations. \BGT\ can automatically simplify these equations; this is discussed in Sect. \ref{sec:reductions}. A list of basic components can be found in Table~\ref{tab:equivalent_models}.

% The functionality of \lstinline{new} command was built on a future-proofing assumption; that users, both human and other software, are likely to want to develop their own component libraries.
% Details of the library interface are beyond the scope of this work, interested readers are invited to refer to the API documentation.

% float 1: list of some common components
% conservation laws, resistance, capacitance, inductance, transformers, gyrators, sources and port 
\begin{table*}
    \centering
	\footnotesize
	\caption{\label{tab:equivalent_models}Basic bond graph variables and components, and their analogues in different physical domains.}
	\renewcommand{\arraystretch}{1.2}
%	\begin{tabular}{c c c c c}
%		\toprule
%		\textbf{Bond graph} & \textbf{Chemical}  & %\textbf{Electrical} & \textbf{Mechanical} & \textbf{Hydraulic}\\ %\midrule
%		\multirow{2}{*}{Effort ($e$)} & Chemical potential & %Voltage & Force & Pressure \\
%		 & $\mu$ [J/mol] & $V$ [V] & $F$ [N] & $P$ [Pa] \\[0.2cm]
%		\multirow{2}{*}{Flow ($f$)} & Molar flow rate & Current & Velocity & Volume flow \\
%		 & $v$ [mol/s] & $I$ [A] & $v$ [m/s] & $Q$ [$\si{m^3/s}$] \\[0.2cm]
%		\textbf{C} & Chemical species & Capacitor & Spring & Tank  \\
%		\textbf{R} & Reaction & Resistor & Damper & Pipe \\
%		\textbf{Se} & Chemostat & Voltage source & Applied force & Pressure source \\
%		\textbf{TF} & Stoichiometry & Transformer & Lever & Pressure transformer \\
%		\textbf{0} & Common species & Parallel connection & Series connection & Parallel connection \\
%		\textbf{1} & Reactant/product complexes & Series connection & Parallel connection & Series connection \\
%		\bottomrule
%	\end{tabular}\\
%	
		\begin{tabular}{c c c c c}
		\toprule
		\textbf{Bond graph} & \textbf{Chemical}  & \textbf{Electrical} & \textbf{Mechanical} & \textbf{Hydraulic}\\ \midrule
		Potential ($e$) & Chemical potential $\mu$ [J/mol] & Voltage $V$ [V] & Force $F$ [N] & Pressure $P$ [Pa] \\
		Flow ($f$) & Molar flow rate $v$ [mol/s] & Current $I$ [A] & Velocity $v$ [m/s] & Volume flow $Q$ [$\si{m^3/s}$]\\
		\textbf{C} & Chemical species & Capacitor & Spring & Tank  \\
		\textbf{R} & Reaction & Resistor & Damper & Pipe \\
		\textbf{Se} & Chemostat & Voltage source & Applied force & Pressure source \\
		\textbf{TF} & Stoichiometry & Transformer & Lever & Pressure transformer \\
		\textbf{0} & Common species & Parallel connection & Series connection & Parallel connection \\
		\textbf{1} & Reactant/product complexes & Series connection & Parallel connection & Series connection \\
		\bottomrule
	\end{tabular}\\
%	
%	\caption[Equivalent models in \BGT.]{Bond graph modelling groups physical processes by function (on energy/power) instead of by physical realisation in a particular domain. Here we list a few of the basic models in the bond graph framework and common physical realisations.}
%	\renewcommand{\arraystretch}{1.0}
\end{table*}

In bond graphs, network conservation laws, such as Kirchhoff's Laws, are themselves considered components, and hence must be added just as any other model.
For example, the \zero\ junction describes the conservation law where all efforts are equal. Suppose there are $n$ ports associated with this junction, then the constitutive relation is
\begin{align}
0 = e_k - e_0\quad \forall\ k\in 1, \ldots, n-1, \qquad \sum_{k=0}^{n-1} f_k = 0.
\end{align}
It is clear by inspection that this is indeed power conserving. Similarly, the $n$-port \one\ junction, or `common flow', has relations given by
\begin{align}
0 = \sigma_jf_j - \sigma_0f_0\quad \forall k \in 1, \ldots{n-1},\qquad \sum_{k=0}^{n-1}\sigma_ke_k =0 \label{eq:one_junction}
\end{align}
where $\sigma_k = 1$ if the associated port $k$ is oriented inwards, or $\sigma_k = -1$ if oriented outwards. 
%Explicitly associating the direction with the port (and hence the junction), instead of determining it circumstantially from the bond means that the governing equations are independent of any particular choice of bond orientation.

\section{Bond graph models of biochemical systems}

The components relevant to biochemical systems are as follows. Note that below, the effort variables $e$ represent chemical potentials $\mu$ and the flow variables represent the fluxes $v$.

\begin{itemize}
\item Species are represented using \Ce\ components -- a nonlinear version of the capacitor (\textbf{C} component). They have the constitutive relationships
\begin{subequations}
\begin{align}
    \dot{x} - f &= 0  \label{eq:Ce_dx} \\
    e - RT \ln (Kx) &= 0
\end{align}
\label{eq:Ce_equations}
\end{subequations}
As mentioned previously, this component contains the species thermodynamic parameter $K$.
\item In cases where a species has a constant concentration, it is represented by the alternative \Se\ (``potential source'') component with a constant chemical potential (which is related to concentration). Such a `chemostat' component has the constitutive relationship
\begin{align}
    e = RT \ln (K\bar{x}) = \text{constant}
\end{align}
\emph{i.e.} this is the same as the \Ce\ component, but with the dynamics (Eq. \ref{eq:Ce_dx}) omitted. Since the concentration is constant, we use the notation $\bar{x}$ to indicate that $d\bar{x}/dt = 0$.
\item Reactions are represented using \textbf{Re} components, qhich are equivalent to two-port nonlinear resistors. These have the constitutive relationships
\begin{subequations}
\begin{align}
    f_1 + f_2 &= 0 \\
    f_1 - r (e^{e_1/RT} - e^{e_2/RT}) &= 0 \label{eq:Re_MdD}
\end{align}
\label{eq:Re_equations}
\end{subequations}
where $e_i$ and $f_i$ are the chemical potentials and fluxes associated with the $i$th bond. These equations are parameterised by the parameter $r$, the reaction rate parameter. The power dissipated is ${P = f_1(e_1-e_2) \ge 0}$ because $f_1 > 0 \Leftrightarrow e_1 > e_2$. 
\item In order to incorporate stoichiometry, \zero\ and \one\ junctions are used. In particular, \zero\ junctions describe the involvement of a species in multiple reactions and the \one\ junction describes the involvement of multiple reactants or products in a single reaction. This is discussed in more detail below.
\item When multiple stoichiometries are involved, \textbf{TF} (transformer) components can be used to scale the chemical potentials and fluxes accordingly. These follow the constitutive relations
\begin{align}
    f_1 - nf_2 = 0 \\
    e_2 - ne_1 = 0
\end{align}
where $n$ is a parameter for the stoichiometry.
\end{itemize}

\begin{figure*}
	\centering
	\includegraphics[width=0.9\linewidth]{closed_biochemical_cycle.pdf}
	\caption{Representations of the reaction network $X \rightleftharpoons Y \rightleftharpoons Z \rightleftharpoons X$. (A) Biochemical network; (B) Bond graph with efforts and flows labelled; (C) The same bond graph as in (B), but with chemical potentials and flux variables computed from conservation laws.
		\label{fig:closed_bg}}
\end{figure*}

To illustrate, we will represent the reaction networks discussed in Sect. \ref{sec:thermodynamic_cycles} as bond graphs. The reaction network $\mathrm{X \rightleftharpoons Y \rightleftharpoons Z \rightleftharpoons X}$ (Fig. \ref{fig:closed_bg}A) is represented by the bond graph in Fig. \ref{fig:closed_bg}B, where each bond has been labelled with its potential and flow variable. As illustrated by the labels, each \Ce\ component (blue) has constitutive relations defined by Eq. \ref{eq:Ce_equations} and each \textbf{Re} component (green) has constitutive relations defined by Eq. \ref{eq:Re_equations}. Since each of the species is involved in two reactions, a common potential (\zero; purple) junction is used to equate the chemical potential contributions of the species to each reaction it is involved in, while simultaneously accounting for conservation of mass. When these conservation equations are accounted for, the efforts and flows can be computed in terms of chemical potentials and fluxes; these are shown in Fig. \ref{fig:closed_bg}C, where the chemical potentials and fluxes are defined as per Eqs. \ref{eq:XYZ_mu}--\ref{eq:XYZ_v}. Thus, by using Eq. \ref{eq:Ce_dx}, it is easily seen that we can recover the dynamics defined by the differential equations
\begin{subequations}
\begin{align}
    \dot{x}_X &= r_3 K_Z x_Z - r_3 K_X x_X - r_1 K_X x_X + r_1 K_Y x_Y \\
    \dot{x}_Y &= r_1 K_X x_X - r_1 K_Y x_Y - r_2 K_Y x_Y + r_2 K_Z x_Z \\
    \dot{x}_Z &= r_2 K_Y x_Y - r_2 K_Z x_Z - r_3 K_Z x_Z + r_3 K_X x_X
\end{align} \label{eq:XYZ_ode}
\end{subequations}


We can apply similar principles in defining the dynamics of the reaction network $\mathrm{X + A \rightleftharpoons Y \rightleftharpoons Z \rightleftharpoons X + B}$ (Fig. \ref{fig:open_bg}A), shown in the bond graph in Fig. \ref{fig:open_bg}B. In addition to the components used in the previous example, we also use \Se\ components and \one\ junctions. The \Se\ components (blue) are used for the species A and B due to the fact that we assume that they are chemostats, \textit{i.e.} have constant concentration. In this second example, some reactions have multiple reactants or products. In these cases common flow junctions (\one; purple) are required to equate their flows while also summing their chemical potential contributions to each reaction; for example, that the forward affinity of the reaction r1 is $\mu_X + \mu_A$. Fig. \ref{fig:open_bg}B shows the chemical potential and flux variables associated with each bond, and it can be shown through substuting the relevant chemical potentials into the Marcelin-de Donder equation (Eq. \ref{eq:Re_MdD}) that the dynamics are governed by the differential equations 
\begin{subequations}
\begin{align}
    \dot{x}_X &= r_3 K_Z x_Z - r_3 K_X x_X K_B \bar{x}_B  \nonumber \\ & \quad - r_1 K_X x_X K_A \bar{x}_A + r_1 K_Y x_Y \\
    \dot{x}_Y &= r_1 K_X x_X K_A \bar{x}_A - r_1 K_Y x_Y  \nonumber \\ &\quad - r_2 K_Y x_Y + r_2 K_Z x_Z \\
    \dot{x}_Z &= r_2 K_Y x_Y - r_2 K_Z x_Z  \nonumber \\ &\quad - r_3 K_Z x_Z + r_3 K_X x_X K_B \bar{x}_B
\end{align}
\end{subequations}
which as expected, is identical to the equations for the energy-based approach (Eqs. \ref{eq:mass_conservation},\ref{eq:open_v}).

\begin{figure*}
	\centering
	\includegraphics[width=0.7\linewidth]{open_biochemical_cycle.pdf}
	\caption{Representations of the reaction network $X + A \rightleftharpoons Y \rightleftharpoons Z \rightleftharpoons X + B$. (A) Biochemical network; (B) Bond graph.
		\label{fig:open_bg}}
\end{figure*}

The approach outlined above can be used to build dynamic energy-based models of arbitrary scale, which account for mass, charge and energy conservation. The approach extends naturally to incorporate electrochemical \cite{Pan2018aa} and biomechanical systems. 

\section{BondGraphTools}
\label{sec:bgt}
Broadly speaking, there are two main computational approaches to bond graph modelling; computer aided design tools and mathematical software. Computer Aided Design (CAD) tools follow in the tradition of electrical design automation and include Dymola\footnote{\url{www.3ds.com/products-services/catia/products/dymola/}} and 20-sim\footnote{\url{www.20sim.com}} in which users draft technical schematics of the system from a library of components. Similarly, mathematical software such as MATLAB (Simulink), Maple (MapleSim) or Mathematica (SystemModeller) also allows users to construct graphical representations of the model. While graphical interfaces can be useful in intuitively understanding bond graph models, they lack the capacity to automate the construction of large systems and do not interface very well with the existing ecosystem of numerical analysis software. Furthermore, the vast majority of bond graph software is embedded within proprietory software, limiting the scope for expansion.

%It is worth noting that much of the modelling and simulation tools are built upon, or interact with Modelica\footnote{\url{www.modelica.com}}, a unified object-oriented markup language for systems modelling.
%However, there does not seem to be much scope within the Modelica ecosystem for a scripting interface, and most Modelica compatible software seems predominately focussed on numerical simulation.

%\begin{figure}
%	\label{fig:bondgraph}
%\caption{An acausal bond graph model of a driven parallel capacitor and inductor network under load. Here the capacitor is represented by the 'C', inductor as 'I' and resistive load component as 'R' respectively 
%Parallel law is represented by the '0', or \emph{equal effort junction}. In this case, the system is driven by a current, or flow, source 'Sf'.
%Since bond graphs are domain agnostic, this model could equivalently represent a damped, driven spring and mass, or the analogical hydraulic system.}	
%\end{figure}

% Written in python, which means open-source, 

\BGT\ aims to resolve the above issues by providing a scripting interface rather than a graphical interface, and is developed within the open-source python language. As a programming interface, \BGT\ gives modellers a means to integrate the tools and techniques of software development into their modelling workflow.
This includes being able to script tasks like model re-parametrisation and batch replacement of model subcomponents both of which can be tedious and time-consuming in graphical environments. 

Building upon and integrating with the existing python ecosystem means that \BGT\ can specialise in providing an interface for model building without concerning itself with other tasks. This results in a smaller codebase, and hence more sustainable software.
Using modern open source practices allows other developers to easily modify, contribute and build upon the \BGT\ codebase. Unlike proprietary software, users are free to implement new features and extend \BGT\ as they see fit, for example by using \BGT\ as a foundation for graphical modelling environments.

For a large class of systems, particularly in systems biology, only the network topology of a system may be known at the time of modelling.
As \BGT\ represents model parameters symbolically, values are free to be determined later in the modelling process via existing parameter estimation techniques.

\BGT\ provides a programming language for building bond graph models which are automatically converted into differential equations to be analysed or simulated.
Python is well established as a robust and easy-to-use general purpose programming language with a wide variety of standardised and well supported libraries for standard scientific tasks, and \BGT\ adheres to python language idioms by emphasising self-explanatory, self-documenting and self-contained code.

\section{Design motivation and basic use}\label{sec:basicuse}
As discussed in Sect. \ref{sec:bg_modelling}, bond graphs are defined in terms of their subsystems are and how they are related.
Here we outline the relevant classes of models and how they are related in \BGT.
Throughout the code examples, we assume \BGT\ has been imported as follows:\\
\begin{minipage}{\linewidth}
\begin{lstlisting}
import BondGraphTools as bgt
\end{lstlisting}
\end{minipage}
We demonstrate code as it appears in a python script, which could be equally executed by entering the code into an IPython session or a Jupyter notebook~\cite{Perez2007}.

\subsection{Creating models}
\label{sec:creation}
% how to create new models and components.
Models in \BGT\ are broadly split into two classes: \emph{atomics}, which represent processes that are considered indecomposable and often fundamental; and \emph{composites}, which are assembled using other models in a has-a relationship. Both of these are constructed using the \lstinline{new} function. 

New composites can be constructed using \\
\begin{minipage}{\linewidth}
\begin{lstlisting}
model = bgt.new(name="New Model")
\end{lstlisting}  
\end{minipage}
which results in the variable \lstinline{model} containing a new instance of the \lstinline{BondGraph} class, the composite base class, with no components and with the name `New Model'.
This is identical to creating new instances of the \lstinline{BondGraph} class directly via\\
\begin{minipage}{\linewidth}
\begin{lstlisting}
model = bgt.BondGraph(name="New Model")
\end{lstlisting}
\end{minipage}
which is available for the purposes of providing an object oriented interface.

New atomics can be created in a similar manner by specifying the model class and value. For example, a \Ce\ component (available in the \lstinline{BioChem} library) can be initialised by\\
\begin{minipage}{\linewidth}
\begin{lstlisting}
species = bgt.new("Ce", library="BioChem",
	value={'k':1,'R':8.314,'T':310})
\end{lstlisting}
\end{minipage}
so that each variable now contains new instances of the respective atomics. This particular atomic has three parameters; the thermodynamic constant $K = 1$, the ideal gas constant $R=8.314$ and the temperature $T=310$.

\subsection{Defining relationships}
There are two categories of relationships between models in \BGT; structural and energetic, corresponding to the components and bonds respectively.
Structural relationships describe composition, how a given model can contain many simpler models. This organises systems into a tree-like hierarchy of 
parent-children relations between models.
Energetic relationships describe how the ports of a given set of child models are connected within a given parent model.

\subsubsection{Structural relationship (components)}
\label{sec:heirarchy}
% how to define model->component relationships.
In \BGT, components can be added (removed) from composite models with the \lstinline{add} and \lstinline{remove} functions. For example, adding a species component to the composite model can be achieved with\\
\begin{minipage}{\linewidth}
\begin{lstlisting}
bgt.add(model, species)
\end{lstlisting}
\end{minipage}
where the first argument is the parent model, and the remaining arguments are the intended children or components. 

A file system interface navigates the model hierarchy and is implemented within the \lstinline{ModelBase} class from which all models inherit.
In particular,
\begin{itemize}
	\item \lstinline{model.uri} is a uniform resource identifier (URI) locating that particular model.
	\item \lstinline{model.parent} refers to the parent model if it exists. 
	\item \lstinline{model.root} refers to the top of the model tree.
	\item For composite models, \lstinline{Composite.components} will contain a list of sub-models.
\end{itemize}
Atomics are thus analogous to files, and composites to directories, with the root model analogous to the unix hostname. 

% talk about why this is great.
\subsubsection{Energetic relationships (bonds)}
\label{sec:bonds}
% how to define component->component relationships.
The energetic relationship defines how power is transferred between components inside a particular model via ports which belong to components, and depends on first establishing a model hierarchy.
Once a set of component-wise relationships is established within a model, components are connected to each other (and hence energy bonds defined, in the bond graph terminology).
When it is not ambiguous, for example when connecting a one-port component to a junction with arbitrarily many identical ports, it is sufficient to connect the component directly via\\
\begin{minipage}{\linewidth}
\begin{lstlisting}
bgt.connect(component_1, component_2)
\end{lstlisting}
\end{minipage}

Otherwise one must specify the port by index\\
\begin{minipage}{\linewidth}
\begin{lstlisting}
target_port = (component_2, 0)
\end{lstlisting}
\end{minipage}
and connect the port directly:\\
\begin{minipage}{\linewidth}
\begin{lstlisting}
bgt.connect(component_1, target_port)
\end{lstlisting} 
\end{minipage}

Composite models keep track of the energy relationships (bonds) in the member attribute \lstinline{Composite.bonds}. 
Similarly, two components can be disconnected via the \lstinline{disconnect} method which has an identical interface to \lstinline{connect}.

\subsection{Model attributes}
Composite and Atomic models often have associated parameters which can be accessed by the member attribute \lstinline{ModelBase.params}. In keeping with the emphasis on symbolic equations, parameters can be either numeric or symbolic values. Once the structure of a composite model has been established, and the internal connections defined, one can generate governing equations for the entire model via \lstinline{ModelBase.constitutive_relations} in terms of the derived state variables \lstinline{ModelBase.state_vars} to be used for further analysis.


\section{Symbolic composition and reduction}
\label{sec:reductions}
Once a model has been constructed, simplified equations are automatically derived using symbolic algebraic tools.
Having a simplified symbolic representation of the system is valuable as it provides modellers a way to export the equations into whatever format they desire. 
In particular, the set of implicit equations can easily be fed into standard parameter estimation routines, discretised for implementation in other architectures, or passed to solver routines.

Core to \BGT\ is symbolic model reduction. 
This relies on the fact that each model or component has a set of constitutive relations $\Phi(X) = 0$ which are implicit equations defining the model behaviour in terms of how power is manipulated.
\subsection{Model structure}
The local co-ordinate space of a model $\alpha$ is taken to be $\mathcal{X}_\alpha = \left\{\dot{x}_\alpha, e_\alpha, f_\alpha, x_\alpha, u_\alpha \right\}$ where $x, \dot{x}$ are the vectors of storage co-ordinates and time derivatives respectively, $e,f$ are the power interconnection variables effort and flow, and $u$ are controls or inputs.
Let us assume that 
$\dim{x_\alpha} = n_\alpha$, $\dim{e_\alpha} = \dim{f_\alpha} = m_\alpha$, $\dim{u_\alpha} = k_\alpha$ for finite $n_\alpha,m_\alpha,k_\alpha$ and $n_\alpha + m_\alpha \ge 1$ so that there is non-trivial behaviour.
We also define $N_\alpha = 2(n_\alpha + m_\alpha) +  k_\alpha$. One can define a matrix $L_\alpha\in \mathbb{R}^{N_\alpha \times N_\alpha}$, a vector field $V_\alpha:\mathcal{X_\alpha}\rightarrow \mathbb{R}^{N_\alpha}$ such that the constitutive relations are equivalent to
\begin{align}
\mathbf{0} = L_\alpha X + V_\alpha (X) \qquad \forall X \in \mathcal{X}_\alpha.
\end{align}
We expect $L_\alpha$ to be sparse and $\mathop{\mathrm{rank}}L_\alpha \le n_\alpha + m_\alpha +  k_\alpha < N_\alpha$ so that there is at least one eigenspace of $L_\alpha$ per state variable pair, unconnected port or control input.
\subsection{Composition}
Any number of constitutive relations ${\Phi = [\Phi_\alpha, \Phi_\beta,\ldots] = 0}$ can thus be combined via
\begin{align}
0 &= LX+V(X) \notag\\ &=  \left(
\begin{matrix}
L_\alpha & 0 &\dots \\
0 & L_\beta & 0 \\ 
\vdots&0 & \ddots
\end{matrix}
\right) X + \left(\begin{matrix}V_\alpha\circ \pi_\alpha \\ V_\beta\circ \pi_\beta \\\vdots \end{matrix}\right)(X) 
\label{eq:implicit_eq}
\end{align}
where $X\in \mathcal{X} =  \mathcal{X}_\alpha \oplus \mathcal{X}_\beta \oplus \ldots$ is the direct sum of local co-ordinate spaces and 
$\pi_\alpha:\mathcal{X} \rightarrow \mathcal{X}_\alpha$ is a projection back into local co-ordinates.
One can easily incorporate interconnecting power bonds by noting that a bond connecting port $i$ on component $\alpha$ to port $j$ on component $\beta$ can be represented by the rows
\begin{align}
0 = e_{\alpha;i} - e_{\beta;j}  = \left[\theta_{e;\alpha;i}, -\theta_{e;\beta;j} \right]\left(\begin{matrix}X_\alpha\\ X_\beta\end{matrix}\right)
\end{align}
and
\begin{align}
0 = f_{\alpha;i} + f_{\beta;j}  = \left[\theta_{f;\alpha;i}, \theta_{f;\beta;j} \right]\left(\begin{matrix}X_\alpha\\ X_\beta\end{matrix}\right)
\end{align}
where $\theta_{e;\alpha;i}$ is the co-basis vector such that $\theta_{e;\alpha;i}X_\alpha = e_{\alpha;i}$ i.e,.; $\theta_{e;\alpha;i}$ is a row vector with one in the column corresponding to $e_{\alpha;i}$ (similarly for $\theta_{e;\beta;j}$, and $\theta_{f;\alpha;i}$ and $ \theta_{f;\beta;j}$ for the flow variables).
It follows that the set of all bonds form a junction structure on the larger space $\mathcal{X}$ represented by a full rank matrix $J$ such that $JX = 0$ with row rank identical to the number of bonds.
It is convenient to simply consider this as an additional constitutive relation, and append it to the linearised matrix $L$ in Eq.~\ref{eq:implicit_eq}.

Given the space $X$, there exists an orthonormal permutation matrix $P$ such that $X' = P^{-1}X$, and 
\begin{align}
X' =(\dot{x}_\alpha, \dot{x}_\beta,\ldots e_\alpha, f_\alpha, e_\beta, f_\beta, \ldots, x_\alpha, x_\beta,\ldots, u_\alpha, u_\beta,\ldots).
\end{align}
It also follows from elementary linear algebra that there is exists an invertible matrix $\Lambda$ such that
$L' = \Lambda L P$ is in a reduced upper triangular form (reduced row echelon form with leading terms always on the diagonal), resulting in an exact simplification of Eq.~\ref{eq:implicit_eq}
\begin{align}
\Lambda L X + \Lambda V(X) = L'X' + V'(X') = 0 \label{eq:simplified}
\end{align}
where $V'(X') = \Lambda V(PX')$.
The coordinate ordering, and hence the permutation matrix $P$ is chosen so that (in the linear case) triangularisation produces the correct order of dependence when substitution is performed; rates of change, efforts and flows are expressed in terms of state and control variables. 
This can be trivially extended to simple nonlinear cases as non-zero diagonal entries of $L'$ determine substitution rules, though more complicated systems can produce irreducible algebraic constraints.
\subsection{Output}
Constitutive relations, retrieved via member attribute \lstinline{ModelBase.constitutive_relations} by evaluating Eq.~\ref{eq:simplified}, can be generated for any model at any level of the structural tree, and is performed in a recursive manner to reduce the number of calculations. 
Reduced symbolic models can be passed into a simulation service, which renders Eq.~\ref{eq:simplified} as a function, then initialises and solves the associated initial value problem, for example by

\noindent
\begin{minipage}{\linewidth}
\begin{lstlisting}
u = "sin(t)"                     
t,x = simulate(model, 			 
			   x0=x0,            
			   timespan=[0,100], 
			   control_vars=[u]),
			   dt=0.1            
\end{lstlisting}
\end{minipage}
Numerical integration is provided by the SUNDIALS~\cite{sundials} suite of differential algebraic equation solvers.

\section{Examples}
\label{sec:example}
\subsection{Closed biochemical cycle}
\label{sec:closed_bgt}
Here we demonstrate the basic features of \BGT\ by constructing the the bond graph of the biochemical cycle in Fig. \ref{fig:closed_bg}. This is shown in the snippet below.

\noindent
\begin{minipage}{\linewidth}
\begin{lstlisting}
def biochemical_cycle(name="Cycle"):
    R = 8.314
    T = 310.0

    model = bgt.new(name=name)
    X = bgt.new("Ce",name="X",library="BioChem",
                 value={'k':1,'R':R,'T':T})
    Y = bgt.new("Ce",name="Y",library="BioChem",
                 value={'k':2,'R':R,'T':T})
    Z = bgt.new("Ce",name="Z",library="BioChem",
                 value={'k':3,'R':R,'T':T})
    common_X = bgt.new("0",name="X")
    common_Y = bgt.new("0",name="Y")
    common_Z = bgt.new("0",name="Z")
    r1 = bgt.new("Re",name="r1",library="BioChem",
                 value={'r':1,'R':R,'T':T})
    r2 = bgt.new("Re",name="r2",library="BioChem",
                 value={'r':2,'R':R,'T':T})
    r3 = bgt.new("Re",name="r3",library="BioChem",
                 value={'r':3,'R':R,'T':T})
    bgt.add(model,X,Y,Z,common_X,common_Y,common_Z,
            r1,r2,r3)

    bgt.connect(common_X,X)  # bond 0
    bgt.connect(common_X,r1) # bond 1
    bgt.connect(r1,common_Y) # bond 2
    bgt.connect(common_Y,Y)  # bond 3
    bgt.connect(common_Y,r2) # bond 4
    bgt.connect(r2,common_Z) # bond 5
    bgt.connect(common_Z,Z)  # bond 6
    bgt.connect(common_Z,r3) # bond 7
    bgt.connect(r3,common_X) # bond 8
    
    return model

model1 = biochemical_cycle()
\end{lstlisting}
\end{minipage}

Line 5 instantiates a new model. Lines 6--22 create the components in the bond graph model and add them to the model. Finally, the components are connected in lines 24--32, with the bonds labelled according to their indices in Fig. \ref{fig:closed_bg}B

Note that for this particular model, we define a function \lstinline{biochemical_cycle} to construct the model. One of the advantages offered by implementing \BGT\ in a scripting language is that functions can be used to make multiple copies of the same template. Later, we use this function as a basis for constructing the similar model of an open biochemical cycle (Fig. \ref{fig:open_bg}).

For this model, we use the parameters $(K_X, K_Y, K_Z) = (1,2,3)\ \si{mM^{-1}}$  and $(r_1,r_2,r_3) = (1,2,3)\ \si{s^{-1}}$. These are added to the model by passing in value arguments; \texttt{'k'} refers to the species constant $K$ and \texttt{'r'} refers to the reaction rate $r$. For \Ce\ and \textbf{Re} components, the ideal gas constant $R$ and temperature $T$ also need to be specified.

Once the components, connections and parameters have been defined, \BGT\ will automatically derive the constitutive relations through the command \lstinline{model1.constitutive_relations}. The output is shown on the snippet below

\begin{lstlisting}
[dx_0 + 4*x_0 - 2*x_1 - 9*x_2,
 dx_1 - x_0 + 6*x_1 - 6*x_2,
 dx_2 - 3*x_0 - 4*x_1 + 15*x_2]
\end{lstlisting}
 
As can be seen, these correspond to Eq. \ref{eq:XYZ_ode} once the parameters have been substituted, where the state variables \texttt{x\_0}, \texttt{x\_1} and \texttt{x\_2} have been mapped to $x_X$, $x_Y$ and $x_Z$ respectively.

We run simulations of the model using the initial conditions $(x_X, x_Y, x_Z) = (2,2,2)\ \si{mM}$ over the time span $0 \ \si{s}<t<1 \ \si{s}$. In the code below, we run separate simulations of the model with different values of $K_X$.
\begin{minipage}{\linewidth}
\begin{lstlisting}
x0 = [2.0,2.0,2.0]
tspan = (0,1.0)
K_X_vals = [1.0, 2.0, 3.0, 4.0]
sol_t = []
sol_x = []
for K_X in K_X_vals:
    (model1/"C:X").set_param('k',K_X)
    t,x = bgt.simulate(model1,tspan,x0,dt=0.01)
    sol_t.append(t)
    sol_x.append(x)
\end{lstlisting}
\end{minipage}

We use line 7 to change the value of $K_X$ once the model has been created. Here, the uri is used to retrieve the \Ce:A component (\lstinline{model1/"C:A"}), following which the \lstinline{.set_params} method is used to alter its $K$ value.

\begin{figure}
    \centering
    \includegraphics[width=0.9\linewidth]{biochemical_cycle_sim}
    \caption{Time-dependence of $v_1$ for the reaction network ${X \rightleftharpoons Y \rightleftharpoons Z \rightleftharpoons X}$.}
    \label{fig:cycle_sim}
\end{figure}

The flux through reaction 1 (Eq. \ref{eq:v1}) is plotted against time in Fig \ref{fig:cycle_sim}. As can be seen, regardless of the value of $K_X$, the flux settles to zero at steady state, as expected of a thermodynamically consistent model. It can easily be verified by modifying the above code that the same is true when the reaction parameters are altered. This illustrates that parameters in an energy-based model can be independently perturbed without violating thermodynamic consistency, which is not the case in general for kinetic models.

\subsection{Open biochemical cycle}
\label{sec:open_bgt}
We next create a bond graph model of the open biochemical cycle in Fig. \ref{fig:open_bg} using \BGT. While we could use the same approach as in above section, here we demonstrate the flexibility of \BGT\ by instead making an incremental change to the above model. This is demonstrated in the code snippet below.

\noindent
\begin{minipage}{\linewidth}
\begin{lstlisting}
model2 = biochemical_cycle(name="Open cycle")
r1 = model2/("R:r1")
r3 = model2/("R:r3")
common_X = model2/("0:X")
bgt.disconnect(common_X,r1)
bgt.disconnect(r3,common_X)

R = 8.314
T = 310.0
K_A = 1
x_A = 1
K_B = 2
x_B = 1

A = bgt.new("Se",name="A",
            value={'e':R*T*np.log(K_A*x_A)})
B = bgt.new("Se",name="B",
            value={'e':R*T*np.log(K_B*x_B)})
XA = bgt.new("1",name="XA")
XB = bgt.new("1",name="XB")
bgt.add(model2,A,B,XA,XB)

bgt.connect(common_X,XA)
bgt.connect(A,XA)
bgt.connect(XA,r1)
bgt.connect(r3,XB)
bgt.connect(XB,common_X)
bgt.connect(XB,B)
\end{lstlisting}
\end{minipage}

We first create a new copy of the closed biochemical cycle and assign it to the \texttt{model2} variable (line 1). Following this, in lines 2--6 we use the uri interface to retrieve and disconnect the connections of the \zero\ junction connected to \Ce:X from the reactions \textbf{Re}:r1 and \textbf{Re}:r2. Finally, we add two \Se\ components and two \one\ junctions to the model and then connect them according to Fig. \ref{fig:open_bg}. By default, the parameters have been set to $K_A=1\ \si {mM^{-1}}$, $K_B=2\ \si {mM^{-1}}$, $\bar{x}_A=\bar{x}_B=1\ \si {mM}$.

\begin{figure}
    \centering
    \includegraphics[width=0.9\linewidth]{open_biochemical_cycle_sim}
    \caption{Simulations of the reaction network $X + A \rightleftharpoons Y \rightleftharpoons Z \rightleftharpoons X + B$. The affinities have been plotted against the steady-state cycle fluxes.}
    \label{fig:open_cycle_sim}
\end{figure}

We next run simulations of the model for different values of $\bar{x}_A$, varying from 0.02 to 200 $\si{mM}$ (Fig. \ref{fig:open_cycle_sim}). The simulations are run from 0 to 10 seconds to ensure the model reaches a steady state, and for each simulation, the flux $v_1$ is recorded at $t = 10\ \si{s}$ and plotted against the cycle affinity $A = \mu_A - \mu_B$. As expected, the steady-state flux has the same sign as the affinity. The cycle has the overall reaction $\mathrm{A \rightleftharpoons B}$, and the laws of thermodynamics dictate that the conversion of A to B will only proceed if $\mu_A > \mu_B$, regardless of the complexity of the transition. This is reflected in the plot being constrained to the bottom left and upper right quadrants. Similarly, the cycle stalls at $\bar{x}_A = 2\ \si{mM}$, where $\mu_A = \mu_B = RT \ln (2)$, which also corresponds to the equilibrium condition $x_B/x_A = K_A/K_B = 1/2$. These results are an illustration of the fact that energy-based models ensure open cycles are consistent with the laws of thermodynamics, just as they are with closed cycles.

\subsection{Multisite phosphorylation}
To further demonstrate the the capabilities that \linebreak \BGT\ has for automation, we make a model of a more complex multisite phosphorylation system. As discussed in Mallavarapu \textit{et al.} \cite{Mallavarapu2008Programming}, the phosphorylation of proteins is an interesting case study for modular modelling approaches as these networks can be highly complex and are capable of reaching multiple steady states. Protein phosphorylation can have three types of properties:
\begin{enumerate}
\item The number of sites available for phosphorylation
\item They can be \textit{sequential} or \textit{non-sequential}. Sequential phosphorylation is when sites must be phosphorylated in a specific order, whereas sites in non-sequential phosphorylation may be phosphorylated in any order.
\item They can be \textit{distributive} or \textit{processive}. Distributive phosphorylation only allows a single site to be phosphorylated at a time, whereas processive phosphorylation allows multiple sites (up to an integer $p$) to be phosphorylated or dephosphorylated at once. We call $p$ the processivity number, and $p = 1$ for distributive phosphorylation.
\end{enumerate}

\begin{figure*}
    \centering
    \includegraphics[width=0.9\linewidth]{4site_phosphorylation}
    \caption{A reaction scheme of distributive sequential phosphorylation of four sites. The names of each reaction are labelled in blue.}
    \label{fig:4site_phosphorylation}
\end{figure*}

Here we construct a bond graph model of a distributive, sequential four-site phosphorylation network, based on the system described by Thomson and Gunawardena \cite{Thomson2009Multistability}. The reactions for this network are shown in Fig. \ref{fig:4site_phosphorylation}. The substrate is denoted as S, with subscripts indicating the number of phosphorylated sites. The kinase is denoted by E and the phosphatase is denoted by F. We have made the following adaptations to the Thomson and Gunawardena model \cite{Thomson2009Multistability} to make it compatible with the energy-based approach:
\begin{enumerate}
\item Since thermodynamic consistency requires reversible reactions, all reactions are assumed to operate in both directions. In cases where reactions were originally defined to be irreversible, the parameters were chosen to give small reverse rate constants.
\item Because energy from the hydrolysis of ATP is required to drive kinases and phosphatases, ATP, ADP and Pi have been added as reactants and products.
\item The energetic parameters $r$ and $K$ were used rather than kinetic parameters $k_i^\pm$
\end{enumerate}
Details of how the parameters were determined are given in \ref{sec:parameters}.

In many cases, there are proteins with as many as 40 phosphorylation sites \cite{Mallavarapu2008Programming}, resulting in complex phosphorylation mechanisms. Automation by computer software can significantly reduce the time to develop models of these systems and reduce the occurence of errors. Here we introduce an an alternative method of constructing biochemical models available in \BGT: the reaction builder. In the code snippet below, we show how a series of reactions can be defined as a series of strings, with a bond graph model algorithmically constructed from the reaction network in Fig. \ref{fig:4site_phosphorylation}.

\noindent
\begin{minipage}{\linewidth}
\begin{lstlisting}
from BondGraphTools.reaction_builder \
	import Reaction_Network
rn = Reaction_Network(
    name="4-site sequential phosphorylation")
rn.add_reaction("E + S0 + ATP = ES0",name="e1a")
rn.add_reaction("ES0 = E + S1 + ADP",name="e1b")
rn.add_reaction("F + S1 = FS1",name="f1a")
rn.add_reaction("FS1 = F + S0 + Pi",name="f1b")
rn.add_reaction("E + S1 + ATP = ES1",name="e2a")
rn.add_reaction("ES1 = E + S2 + ADP",name="e2b")
rn.add_reaction("F + S2 = FS2",name="f2a")
rn.add_reaction("FS2 = F + S1 + Pi",name="f2b")
rn.add_reaction("E + S2 + ATP = ES2",name="e3a")
rn.add_reaction("ES2 = E + S3 + ADP",name="e3b")
rn.add_reaction("F + S3 = FS3",name="f3a")
rn.add_reaction("FS3 = F + S2 + Pi",name="f3b")
rn.add_reaction("E + S3 + ATP = ES3",name="e4a")
rn.add_reaction("ES3 = E + S4 + ADP",name="e4b")
rn.add_reaction("F + S4 = FS4",name="f4a")
rn.add_reaction("FS4 = F + S3 + Pi",name="f4b")

rn.add_chemostat("ATP")
rn.add_chemostat("ADP")
rn.add_chemostat("Pi")

model = rn.as_network_model(normalised=True)
\end{lstlisting}
\end{minipage}

As illustrated by the code, the \lstinline{reaction_builder} module has the following functionalities:
\begin{enumerate}
\item A new reaction network can be initialised using the \lstinline{Reaction_Network} class (lines 3--4)
\item New reactions can be added using the \lstinline{.add_reaction} method (lines 5--20)
\item Chemostats can be labelled using the \lstinline{.add_chemostat} method. (lines 22--24). In this example, we assume that the concentrations of ATP, ADP and Pi are maintained at constant levels, and thus define them as chemostats. 
\item A bond graph model can be algorithmically built from the reaction network using the \lstinline{.as_network_model} method (line 26).
\end{enumerate}

Once the bond graph model has been constructed, the usual functions and methods in \BGT\ can be used to define parameters and run simulations (see Sects. \ref{sec:closed_bgt}--\ref{sec:open_bgt}).

We simulate the model with the initial conditions $[E] = [F] = 2.8\ \si{\mu M}$, $[S_0] = \alpha S_\text{tot}$ and $[S_4] = (1-\alpha) S_\text{tot}$, where $\alpha$ is a parameter varied between 0 and 0.98 and $S_\text{tot} = 10\ \si{\mu M}$. The simulation results in Fig. \ref{fig:multisite_simulation} show that the system exhibits three distinct steady states (lines in blue, green and red), which is consistent with previous results on these types of models \cite{Mallavarapu2008Programming,Thomson2009Multistability}.

\begin{figure}
    \centering
    \includegraphics[width=0.9\linewidth]{multisite_phosphorylation_sim}
    \caption{Simulations of the distributive sequential four-site phosphorylation model. The simulations were run with the initial conditions $[E] = [F] = 2.8\ \si{\mu M}$, $[S_0] = \alpha S_\text{tot}$ and $[S_4] = (1-\alpha) S_\text{tot}$, where $\alpha$ is varied between 0 and 0.98 and $S_\text{tot} = 10\ \si{\mu M}$. All other concentrations had an initial concentration of zero. Depending on the initial conditions, the system can achieve one of three steady states; these are grouped into blue, green and red lines.}
    \label{fig:multisite_simulation}
\end{figure}

We note that since \BGT\ is embedded in python, it is very flexible in generating models of with similar but non-identical structures. For example, the code for this example could easily be generalised to construct energy-based models of distributive phosphorylation with an arbitrary number of sites $n$ and processivity $p$. This is shown in the code snippet below, where the function \lstinline{multisite(n,p)} will create a bond graph model of a phosphorylation system with $n$ sites and processivity $p$.

\noindent
\begin{minipage}{\linewidth}
\begin{lstlisting}
def multisite(n_sites,processivity=1,
              name="Multisite Phosphorylation"):
    substrates = [f"S{i}" for i in range(n_sites+1)]
    (E,F,ATP,ADP,Pi) = ("E","F","ATP","ADP","Pi")

    rn = Reaction_Network(name=name)
    for p in range(1,processivity+1):
        for i in range(n_sites+1-p):
            Sa = substrates[i]
            Sb = substrates[i+p]
            if p == 1:
                p_str = ""
            else:
                p_str = f"{p}*"
                
            # Kinase reactions
            rn.add_reaction(
                f"{Sa}+{E}+{p_str}{ATP}="+
                f"{E}{Sa}_{p_str[:-1]}{ATP}",
                name=f"e({i},{i+p})a")
            rn.add_reaction(
                f"{E}{Sa}_{p_str[:-1]}{ATP}="
                +f"{Sb}+{E}+{p_str}{ADP}",
                name=f"e({i},{i+p})b")
            
            # Phosphatase reactions
            rn.add_reaction(
                f"{Sb}+{F}={F}{Sb}",
                name=f"f({i+p},{i})a")
            rn.add_reaction(
                f"{F}{Sb}={Sa}+{F}+{p_str}{Pi}",
                name=f"f({i+p},{i})b")

    rn.add_chemostat(ATP)
    rn.add_chemostat(ADP)
    rn.add_chemostat(Pi)

    model = rn.as_network_model(normalised=True)
    return model
\end{lstlisting}
\end{minipage}

\section{Discussion}
\label{sec:discussion}
Biological systems are constrained by the laws of physics in general and the laws of thermodynamics in particular. Here we have argued for the use of an energy-based approach for developing models consistent with the laws of physics. Bond graphs are an established modelling tool for enabling this approach, and we have introduced  \BGT, a software package that adds a layer of automation to bond graph modelling.

\subsection{Model abstraction}
The complexity of large projects, modelling or otherwise, is often made tractable by introducing abstractions which allow individuals to `divide-and-conquer' the problem. For this to work, one requires a way for subcomponents to interface with each other. Bond graphs bring this to modelling by framing the interconnection in terms of power, which in turn allows \BGT\ to define a consistent way of joining sub-models together (via ports). Hence, one avoids some connectivity issues such an inability for models to interface and failing to account for loading, which are issues that commonly arise in systems and synthetic biology \cite{Szigeti2018Blueprint,DelVecchio2013Control}. Furthermore, the energy-based parameterisation of biochemical systems that an energy-based approach provides has the potential to resolve potential inconsistencies between models \cite{Mason2019Energetic}.

Over recent years, systems biologists have favoured a ``white-box" approach to modularity. Here, each module is itself a simulatable model that can be later merged or composed with other models, possibly through semantic annotation of components \cite{Neal2014Reappraisal,Neal2015Semantics}. White-box modularity requires connections to be exposed on the fly and implementing this approach requires a degree of flexibility that is unavailable through traditional graphical interfaces for bond graph software. This white-box approach has recently been applied in the context of bond graphs \cite{Gawthrop2017Chemiosmotic,Shahidi2021Semantic}, and \BGT\ is particularly suited to implementing white-box modularity by being embedded in a scripting environment. Furthermore, python contains an object-oriented interface that could potentially allow an annotation scheme to be built on top of components in \BGT.

\BGT\ has a similar philosophy to the programmatic modelling approach in systems biology, as implemented by \textit{little b} \cite{Mallavarapu2008Programming} and more recently PySB \cite{Lopez2013Programming}. Key features that these packages share are that: (a) each model is an object in a computer program that can be modified, and (b) that they are embedded within a scripting environment, which allows models to be created through highly abstracted means such as functions, rules and macros. However, \BGT\ differs from the above two packages in that it creates energy-based models with energetic parameters, and that it is a general purpose representation, making it more suitable for multi-physical systems such as electrophysiology and mechanochemistry. Further work will aim to provide a richer set of interfaces to \BGT\ for systems biology, including incorporating abstracted approaches such as rule-based modelling \cite{Chylek2014Rule} and building a library of macros for common motifs such as enzyme-catalysed reactions.

\subsection{Towards multi-physics, multiscale models}
\label{sec:multiphysics}
A characteristic of biology is that it spans across multiple physical domains: electrophysiology and redox reactions are electrochemical systems, whereas muscle contraction and mechanosensation are mechanochemical processes. Bond graphs have the potential to drive theoretical advances in these fields due to their generalised nature, which naturally allows models to span across different physical domains. Connections between ports representing different physical domains are made using the energy-transmitting \textbf{TF} (transformer) component~\cite{Gawthrop2007aa,KarMarRos12}. Thus, for example, the chemical and electrical domains are connected using a transformer with modulus $F$; the Faraday constant. Indeed, bond graphs have already been shown to be useful in these areas~\cite{Gawthrop2020Physically,Pan2018aa,Pan2018ab,Gawthrop2020Bioelectricity,Diaz2011Multiscale}. Due to the inherent modularity of bond graphs, such models could potentially be linked models of circulation to create multi-scale models of organs \cite{deBono2018Multiscale}.

An area of active research is the construction of fully integrated models of heart cells, incorporating electrophysiology, calcium signalling, contraction and metabolism \cite{Crampin2020Energetics,Tran2015Regulation}. In some cases, heart disease has been hypothesised to arise from the dysregulation of matching energy production to demand. Thus, bond graph models are ideally suited to answering these questions with their ability to model across multiple physical domains and their thermodynamic consistency.

\subsection{Version controlled models}
Because new biological datasets are actively being generated, models in systems biology are frequently updated in light of new measurements, and later models will often inherit parameters and equations from earlier models. It is therefore of critical importance the provenance of models to be recorded, particularly with respect to the relationship between parameters and equations to experimental protocols and data \cite{Daly2018Reproducible}.

In \BGT, models are constructed via an API, and hence the entire process is textual. 
This means that models can be version controlled using standard concurrent versioning systems (CVS) such as git, mercurial or svn.  In Sect. \ref{sec:open_bgt}, we discussed one way in which \BGT\ could be relevant in this space by reusing a template to make an incremental change to a model. We envision that libraries of models created by \BGT\ could be assembled, shared and updated in the same way one would package and distribute a python script. While the application of \BGT\ here is a subject of future work, it is clear that the utility of integrating efficient and well established `off-the-shelf' version tracking along with the distribution of model libraries cannot be understated.

\subsection{Sustainable software practices}\label{sec:sussoft}
The development of \BGT\ employs many sustainable software techniques which benefit both developers and users.
% Package Availablity and backwards compatability: PyPI 
\BGT\ is available on the python package index (PyPI) and requires python 3.7. 
The latest version can be installed using the console command \texttt{pip3 install BondGraphTools} and requires only the python and SUNDIALS binaries to be pre-installed.
% Documentation: readthedocs
Further installation guides, tutorial, and code documentation is hosted on readthedocs\footnote{\url{http://bondgraphtools.readthedocs.io}} which is updated automatically as new versions are released.
% Source version control: github
Source code is accessible online\footnote{\url{https://github.com/BondGraphTools}} and is distributed under the permissive, open-source Apache 2.0 license. 
Semantic versioning via git tags is used to keep track of releases in perpetuity. The code required to run the examples in this paper are available both on GitHub\footnote{\url{https://github.com/uomsystemsbiology/BGT-Biology}} and Zenodo\footnote{\url{https://doi.org/10.5281/zenodo.4626922}} for posterity to capture both the library and dependencies' state at the time of publication to ensure reproducibility~\cite{hurley_virtual_2015}.

Development of \BGT\ proceeds within an agile iterative and adaptative methodology, as opposed to using a sequential plan-build-test process, which ensures that usable software is prioritised and 
time is not wasted on planning features that users are less interested in. Instead, requirements and design emerge out of the development process and user experiences, with punctuated breaks for refactoring to simplify the codebase.
Test-driven development (TDD) is employed to ensure quality, continuity and stability with automated integration/testing and code metrics provided via Travis.ci\footnote{\url{https://www.travis-ci.com}} and CodeClimate\footnote{\url{www.codeclimate.com}} respectively.
With TDD, unit tests for new software features are specified and implemented before development begins. A feature is then complete once it passes the corresponding unit tests (in addition to the existing tests) prompting integration of the new code into the trunk of code base. TDD is eminently suited to API or library development as developing a battery of tests for common use cases is an integral part of the development process ensuring that existing functionality does not break as new features are added and encouraging developers to think hard about the library interface.
At the time of writing, test coverage is around 80\%, which is respectable in the context of an active, iterative  development process.
 
The feature development and bug status can be tracked using the GitHub issue interface. Users are encourage to log bug reports about any issues they might have.

\section{Conclusion} \label{sec:conc}
Specialist tools are required to build and analyse complex biophysical systems and these often take the form of monolithic computer aided design applications, or extensions to mathematical software.
Here we have presented \BGT, an open source python library for systems modelling which enables the model building process to be scripted, allowing for automation previously unavailable. 
Further, \BGT\ uses best-practice software engineering techniques to increase the sustainability and longevity of the software.
We have demonstrated how \BGT\ can contribute to the modeller's workflow by considering a number of biological models in the existing literature.
\BGT\ continues to be actively developed and used across a variety of problem domains in systems biology, bio-electrics and electrical engineering, and also has potential applications in optomechanics \cite{Cudmore2015Feb}.
Although basic symbolic model order reduction exists within \BGT, an important future research direction involves developing and implementing scalable symbolic order reduction algorithms for nonlinear systems.
Reduced order equations are a desired output as they are a simpler representation of the system dynamics, giving greater insight into how parameters effect emergent states and speeding up numerical simulations.
This is particularly important for systems with multiple time scales, occurring commonly in biology and engineering, which can pose significant challenges to numerical solvers.
\BGT\ offers an promising platform on which to develop both exact and approximate symbolic reduction algorithms which would be a useful in systems biology, but also more broadly in engineering, scientific computing and computational mathematics.


\begin{acknowledgements}
This research was in part conducted and funded by the Australian Research Council Centre of Excellence in Convergent Bio-Nano Science and Technology (project number CE140100036). PJG would like to thank the Faculty of Engineering and Information Technology, University of Melbourne, for its support via a Professorial Fellowship.
\end{acknowledgements}

\section*{Author contribution statement}
PC designed and wrote the code for \BGT, including developing the methodology for symbolic reduction. MP applied the methodology to biochemical systems. PC and MP wrote the manuscript with support from PJG and EJC. All authors revised and edited the manuscript.

\appendix
\section{Parameters for multisite phosphorylation}
\label{sec:parameters}
When only kinetic data are available, the energetic parameters cannot be determined uniquely due to the relative nature of chemical potential and the absence of reference values. This parameter undeterminacy has been discussed in previous papers, where sets in parameter space with equivalent kinetic behaviours were defined \cite{Gawthrop2015Hierarchical,Pan2018aa,Mason2019Energetic}. However, for this model, we will avoid the issue of parameter uncertainty for simplicity and refer the reader to the above articles for further information.

We make the following assumptions on the energetic parameters:
\begin{itemize}
\item We set $\mu_\text{ATP} = 50\ \si{kJ/mol}$, $\mu_\text{ADP} = 0\ \si{kJ/mol}$ and $\mu_\text{Pi} = 0\ \si{kJ/mol}$ since the free energy of ATP hydrolysis $\Delta G = \mu_\text{ADP}+\mu_\text{Pi}-\mu_\text{ATP}$ typically varies between $-50$ kJ/mol and $-60$ kJ/mol in physiological contexts.
\item We set $K_E = K_F = K_{S_0} = 1\ \si{nM^{-1}}$ (corresponding to a free energy of formation of zero at a standard concentration of 1 nM).
\item We assume that the energetics of phosphorylation are identical for each site, \textit{i.e.} $K_{S_{i+1}} = \gamma K_{S_{i}}$. Hence, in conjunction with the above assumption, $K_{S_i} = \gamma^i$. For reasons that we justify later, we have chosen $\gamma = 3.47 \times 10^4$.
\end{itemize}

Note that implicit in the above assumptions are that ADP, Pi, $\mathrm{S}_0$, E and F are the ``reference species" whose chemical potentials are used to define the potentials of the rest of the species. It is relatively straightforward to adapt the assumptions when the more standard chemical free energies of formation are used, in which the chemical potentials of each species are referenced to their constituent elements in their standard states.

The rest of the parameters can be determined from the kinetic parameters in Thomson and Gunawardena \cite{Thomson2009Multistability}. Omitting chemostats, each of the enzyme-catalysed reactions follows the generic structure
\begin{align}
    \mathrm{S}_u + \mathrm{X} \mathrel{\mathop{\rightleftharpoons}^{a}_{b}} \mathrm{XS}_u \mathrel{\mathop{\rightleftharpoons}^{c}_{d}} \mathrm{S}_v + \mathrm{X}
\end{align}
where $\mathrm{S}_u$ and $\mathrm{S}_v$ are the input substrates, $\mathrm{X = E} \text{ or } \mathrm{F}$ and $\mathrm{XS}_u$ is the complex. The kinetic constants of Thomson and Gunawardena are given in Table \ref{tab:multisite_kinetics}. Since the reaction scheme does not explicitly account for chemostats, they have been absorbed into the kinetic parameters.

We need to determine the rate constants of both reactions and the species constant of $\mathrm{XS}_u$. These can be determined using the following equations, which are derived by writing kinetic parameters in terms of the energetic parameters:
\begin{align}
    r_a &= a/(K_{S_u} K_E e^{A_\text{cs}^f/RT}) \\
    K_{\mathrm{XS}_u} &= b/r_a \\
    r_b &= c/K_{\mathrm{XS}_u}
\end{align}where $A_\text{cs}^f$ and is the potential of the reactant chemostat (if present). Similarly, we can define $A_\text{cs}^r$ as the potential of the product chemostat. Thus, for kinases (E), $A_\text{cs}^f = \mu_\text{ATP}$ and $A_\text{cs}^r = \mu_\text{ADP}$. For phosphatases (F), $A_\text{cs}^f = 0$ and $A_\text{cs}^r = \mu_\text{Pi}$.

The remaining kinetic constant $d = r_b S_v A_\text{cs}^r$ is assumed to be zero in Thomson and Gunawardena, but this is impossible in a real system, which requires all reactions to be reversible. Thus, we choose the final parameter $\gamma$ to minimise the magnitude of these rate constants, or more precisely, we minimise their squared sum
\begin{align}
    J = \sum_{i=1}^4 (d_i^E)^2 + \sum_{i=1}^4 (d_i^F)^2
\end{align}
where $d_i^X$ is the parameter $d$ for the $i$th kinase ($\mathrm{X=E}$) or phosphatase ($\mathrm{X=F}$) reaction. A value of ${\gamma = 3.47 \times 10^4}$ will minimise $J$. The full list of energetic parameters is given in Tables \ref{tab:phos_species}--\ref{tab:phos_reactions}.

\begin{table}
\caption{Kinetic parameters used in the Thomson and Gunawardena model\cite{Thomson2009Multistability}.}
\begin{tabular}{l c c c c}
\toprule
          & $i=1$   &  $i=2$  &  $i=3$  &  $i=4$ \\ \midrule
$a_i^E$ (nM/s) & 0.00812 & 0.102   & 0.00812 & 0.102 \\
$b_i^E$ (nM/s) & 0.016   & 0.204   & 0.016   & 0.204 \\
$c_i^E$ ($\si{s^{-1}}$)  & 0.100   & 10.00   & 0.100   & 10.000 \\ \midrule
$a_i^F$ (nM/s) & 0.112   & 0.00264 & 0.651   & 0.00285\\
$b_i^F$ (nM/s) & 0.224   & 0.005   & 1.30    & 0.006  \\
$c_i^F$ ($\si{s^{-1}}$)  & 11.0    & 0.017   & 63.9    & 0.136 \\ \bottomrule
\end{tabular}
\label{tab:multisite_kinetics}
\end{table}

\begin{table}
\centering
\caption{Species parameters $K$ for the four-site phosphorylation example. All quantities are in $\si{nM}^{-1}$}
\begin{tabular}{c l}
\toprule
Species & Value \\ \midrule
E   & 1.000$\times 10^{0}$ \\ 
F   & 1.000$\times 10^{0}$ \\
$\mathrm{S_0}$  & 1.000$\times 10^{0}$ \\
$\mathrm{S_1}$  & 3.470$\times 10^{4}$ \\
$\mathrm{S_2}$  & 1.204$\times 10^{09}$ \\
$\mathrm{S_3}$  & 4.178$\times 10^{13}$ \\
$\mathrm{S_4}$  & 1.450$\times 10^{18}$ \\
$\mathrm{ES_0}$ & 5.246$\times 10^{8}$ \\
$\mathrm{ES_1}$ & 1.848$\times 10^{13}$ \\
$\mathrm{ES_2}$ & 6.316$\times 10^{17}$ \\
$\mathrm{ES_3}$ & 2.225$\times 10^{22}$ \\
$\mathrm{FS_1}$ & 6.940$\times 10^{4}$ \\
$\mathrm{FS_2}$ & 2.280$\times 10^{9}$ \\
$\mathrm{FS_3}$ & 8.344$\times 10^{13}$ \\
$\mathrm{FS_4}$ & 3.052$\times 10^{18}$ \\ \bottomrule
\end{tabular}
\label{tab:phos_species}
\end{table}

\begin{table}
\centering
\caption{Reaction parameters $r$ for the four-site phosphorylation example. All quantities are in nM/s.}
\begin{tabular}{c l}
\toprule
Reaction & Value \\ \midrule
e1a & 3.050$\times 10^{-11}$ \\
e1b & 1.906$\times 10^{-10}$ \\
f1a & 3.228$\times 10^{-6}$ \\
f1b & 1.585$\times 10^{-4}$ \\
e2a & 1.104$\times 10^{-14}$ \\
e2b & 5.412$\times 10^{-13}$ \\
f2a & 2.193$\times 10^{-12}$ \\
f2b & 7.455$\times 10^{-12}$ \\
e3a & 2.533$\times 10^{-20}$ \\
e3b & 1.583$\times 10^{-19}$ \\
f3a & 1.558$\times 10^{-14}$ \\
f3b & 7.659$\times 10^{-13}$ \\
e4a & 9.170$\times 10^{-24}$ \\
e4b & 4.495$\times 10^{-22}$ \\
f4a & 1.966$\times 10^{-21}$ \\
f4b & 4.456$\times 10^{-20}$ \\ \bottomrule
\end{tabular}
\label{tab:phos_reactions}
\end{table}

% BibTeX users please use one of
%\bibliographystyle{spbasic}      % basic style, author-year citations
%\bibliographystyle{spmpsci}      % mathematics and physical sciences
\bibliographystyle{spphys}       % APS-like style for physics
\bibliography{bibliography.bib}   % name your BibTeX data base

\end{document}
% end of file template.tex
