## Supplementary figures and images for "Analysing and simulating energy-based models in biology using BondGraphTools"

### 4site_phosphorylation.pdf

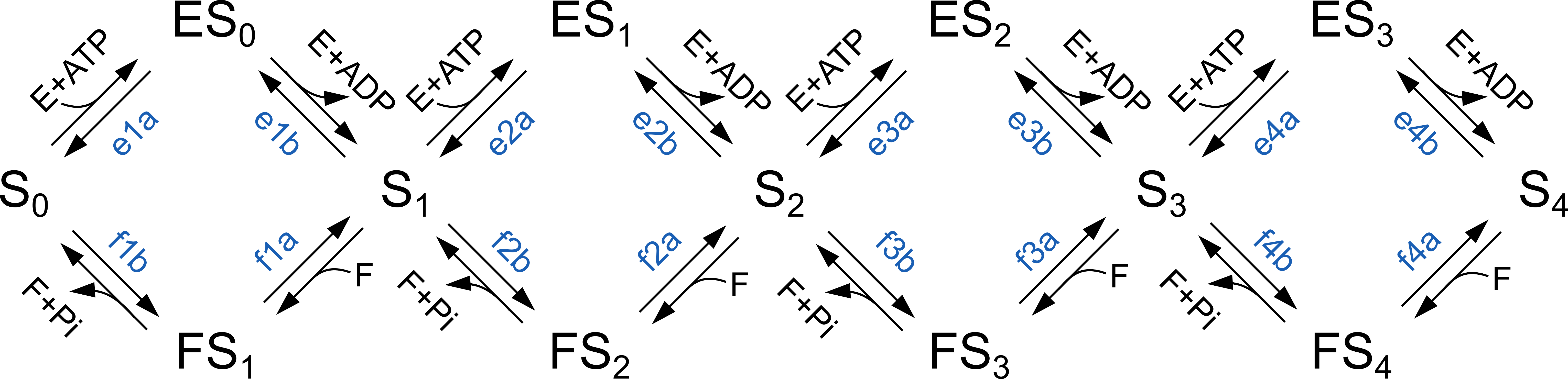

### biochemical_cycle_sim.pdf

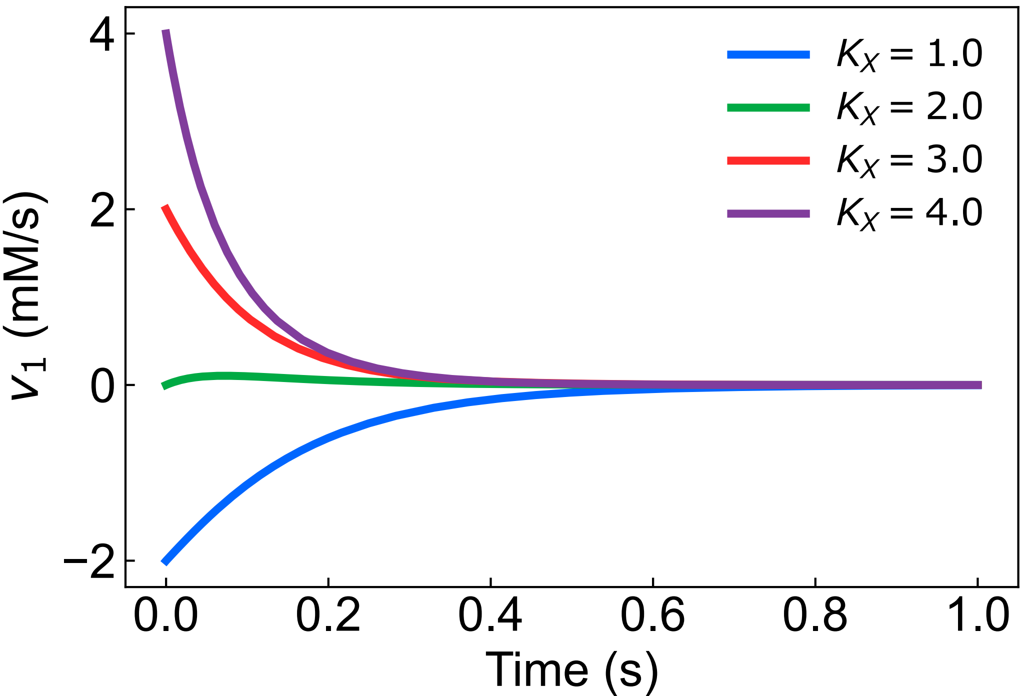

### closed_biochemical_cycle.pdf

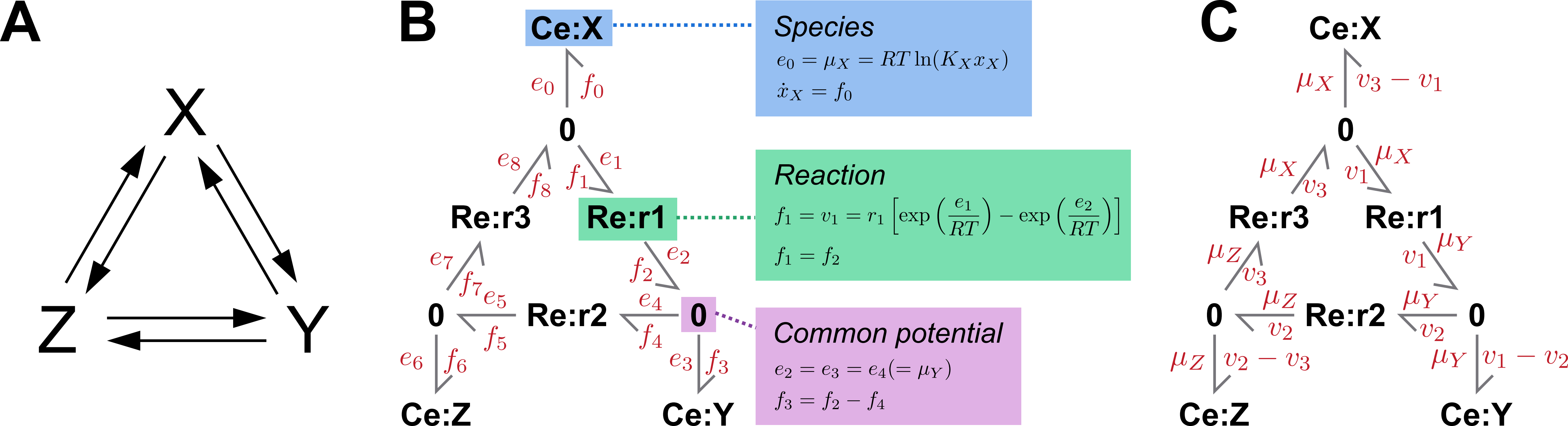

### figure1.pdf

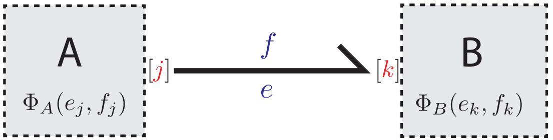

B

$\Phi_B(e_k, f_k)$

### multisite_phosphorylation_sim.pdf

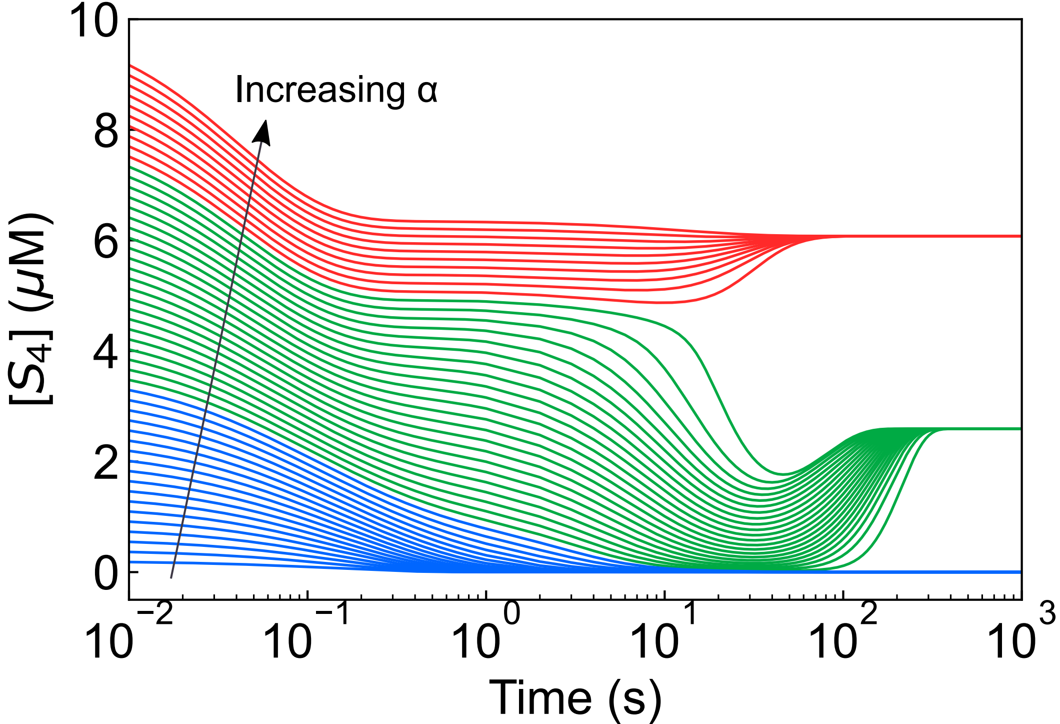

### open_biochemical_cycle.pdf

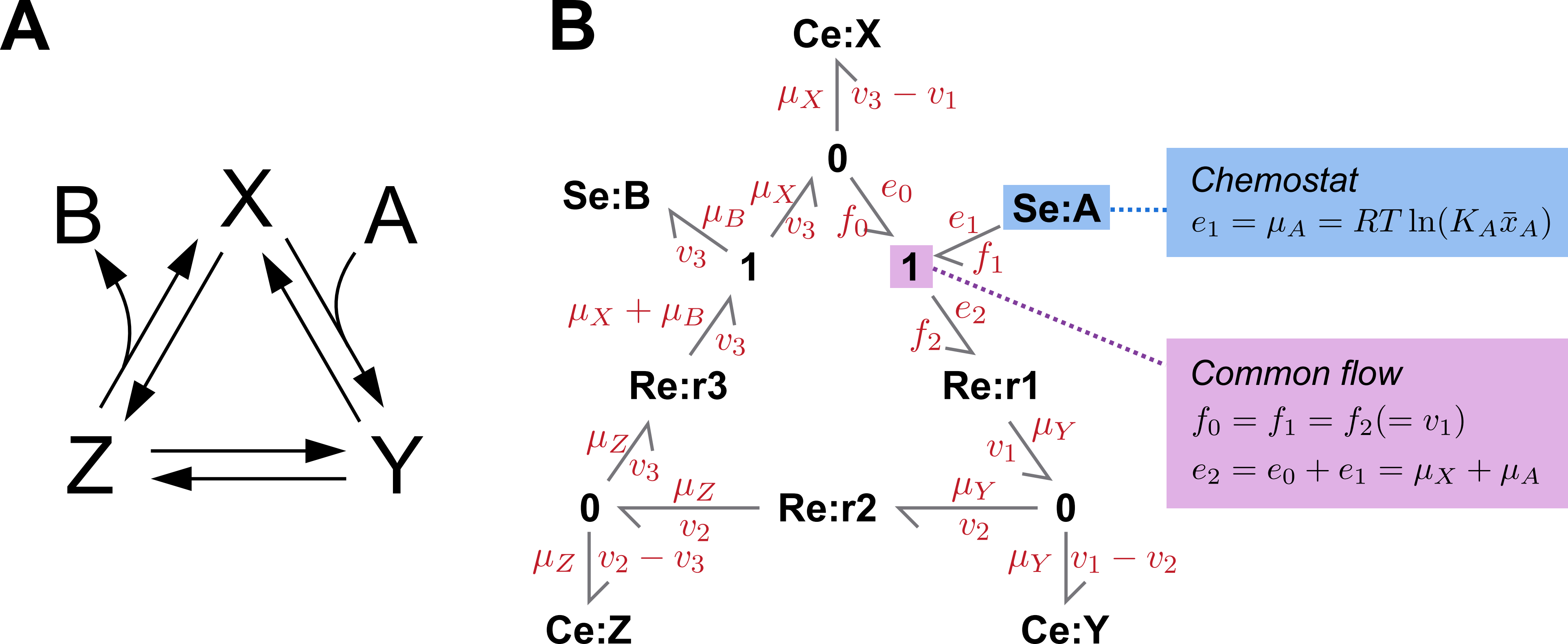

### open_biochemical_cycle_sim.pdf

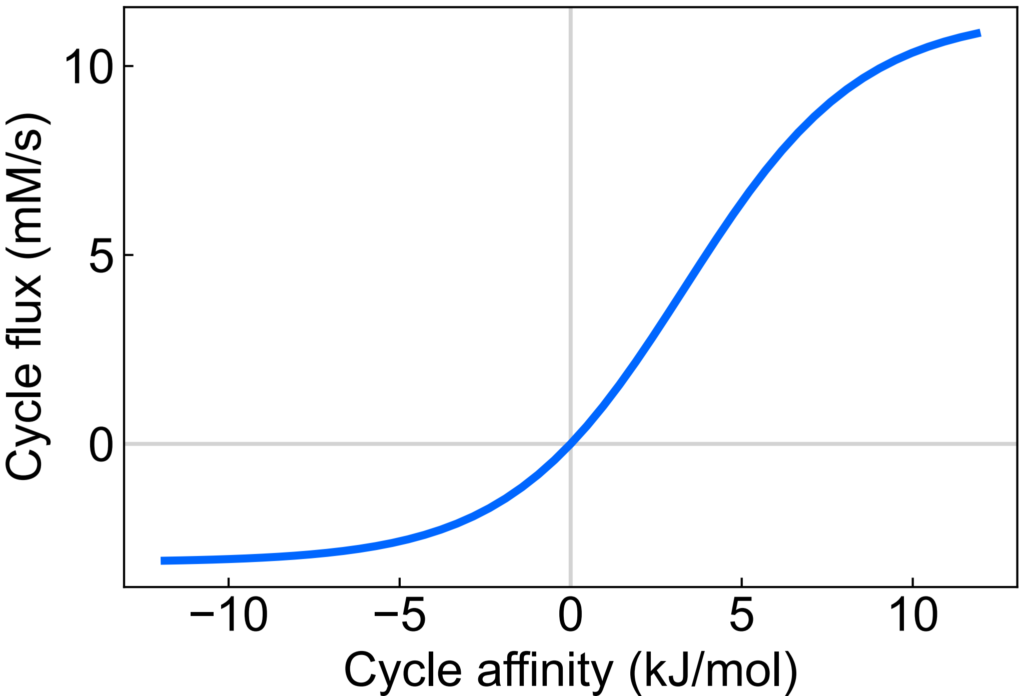
